# The Social Switchboard: Context and Rank Shapes Behavioural and Neural Responses During Macaque Decision-Making

**DOI:** 10.1101/2025.08.11.669617

**Authors:** Shahab Zarei, Jiahao Tu, Xiaochun Wang, Ian Max Andolina

## Abstract

**Key Points:**

- Sociality influences ethologically important decisions and demand a high level of cognitive sophistication
- We used a novel decision making task with 6 macaques where social hierarchy could be tested across social audience, coaction, envy, altruism, cooperation & competition contexts.
- Subject’s decision making performance was significantly modulated by context and social rank, reflected by their social eye gaze patterns.
- There were significant differences in fronto-parietal brain regions as the social context changed

Social interactions shape many of our most ethologically relevant decisions and require considerable cognitive sophistication. Factors such as immediate needs, social bonds, hierarchical status, and reciprocity norms impact decision-making. Yet most laboratory paradigms examine individuals in isolation. We developed a face-to-face cognitive testing platform using a transparent touchscreen that allowed pairs of adult male *Macaca fascicularis* (n = 6) with established dominance relationships to perform a decision-making task across six social contexts: audience, co-action, envy, cooperation, competition, and altruism. We recorded eye movements simultaneously from both subjects (gaze position and pupil diameter), synchronised behavioural video, and task performance. Across social contexts, macaques performed better in the presence of a conspecific than when tested alone, but performance declined under altruistic and competitive conditions. Dominance substantially modulated motivation: dominant subjects were less willing to engage when rewards were shared (envy), when only the partner was rewarded (altruism), or when contingent action with a subordinate was required (cooperation). Eye-tracking revealed greater pre-choice arousal and social attention during social sessions, indexed by larger pupils, longer fixations on the partner’s face, and more saccades before—but not during or after—choices. Pupil dilation was larger in dyads where both individuals were dominant than in dyads where one was subordinate. Finally, noninvasive functional near-infrared spectroscopy (fNIRS) showed increased cortical engagement relative to baseline in audience, altruism, competition, and cooperation conditions. These findings demonstrate that social context and hierarchical dominance systematically shape choices, attentional allocation, autonomic arousal, and cortical engagement in macaques, and provide a tractable platform for further dissecting the neural bases of contextual social decisionmaking.

## 1. Introduction

Many of our most important decisions are influenced by the dynamics of social engagements and the feedback we receive from others. Studies in humans and animals demonstrate that the mere presence of a conspecific alters an individual’s behaviour and the decisions they make (Reynaud *et al*., 2015; Dumontheil *et al*., 2016; Sekiguchi & Hata, 2019). Fundamental behaviours like altruism, envy, and cooperation vary based on many factors, such as need, relationships, authority, power, and reciprocity, highlighting the need to study social decision-making as a complex and complementary approach for understanding cognitive processes. However, how these behaviours are shaped by context and how they impact the underlying neural procession remain poorly understood.

Social interactions require inferring others’ goals, predicting their actions, and responding adaptively. This complexity arises because social decisions not only consider contextual factors from the environment and the stimuli (Zarei *et al*., 2019a, 2019b) but also grapple with the meaning of an action as it varies based on the interaction partner and the agent’s internal state (Stoodley & Tsai, 2021). Evidence from biology and comparative psychology reveals that many animals exhibit a variety of social and prosocial behaviours: parrots cooperate voluntarily for mutual rewards (Brucks & von Bayern, 2020), bonobos share food with companions (Tan & Hare, 2013), and rats help conspecifics that are seeking food (Scheggia & Papaleo, 2020), assist distressed conspecifics (Bartal & Mason, 2018), or reciprocate prior help (Dolivo & Taborsky (2015)). Rodents may console peers (Burkett *et al*., 2016) or cooperate (Choe *et al*., 2017), but their actions are typically driven by immediate social or survival cues rather than higher-order inference, and often lack the cognitive complexity seen in primates, such as altruism, defined by personal cost without immediate benefit (Marshall-Pescini *et al*., 2016), which differs from empathy-driven helping (de Waal, 2008; Clavien & Chapuisat, 2013). These findings underscore the need for studies in non-human primates, whose evolutionary complexity mirrors humans’ social cognition.

Primates in the family *Cercopithecidae* are an ideal model for investigating the neural and cognitive mechanisms of social decision-making due to homologous brain structures and complex social behaviours [e.g., hierarchies, cooperation, and social learning; de Waal (1990); Goodwin (2010)]. Rhesus macaques, for instance, acquire social information through observation (Cheney & Seyfarth, 1990) and can exhibit altruistic behaviours like sacrificing food to alleviate others’ distress (Masserman *et al*., 1964). Their dominance hierarchies and emotional contagion (de Waal & Luttrell, 1985; Nieuwburg *et al*., 2021) further mirror human social dynamics. Critically, their neural architecture (including mirror neuron systems and prefrontal connectivity) closely parallels humans’, enabling translation of findings to social cognition disorders (Brosnan & de Waal, 2003; Rudebeck *et al*., 2006; Kano & Tomonaga, 2009), bridging gaps between basic neuroscience and clinical research (Capitanio & Emborg, 2008; Bauman & Schumann, 2018). This gap is clinically urgent: Social decision-making deficits are implicated in autism (Testard *et al*., 2024), schizophrenia (Green *et al*., 2015), depression (Kupferberg *et al*., 2016), Alzheimer’s (Shi *et al*., 2025), and other psychiatric disorders, with profound health consequences that impact the daily functioning of individuals. The inability to establish meaningful connections with others is closely linked to depression and various long-term detrimental effects on overall health and well-being (Holt-Lunstad *et al*., 2015). Yet the evolutionary and neural bases of many behaviours like altruism, empathy, and envy remain unclear. Ultimately, furthering our basic understanding of the social brain will provide new targets and opportunities to treat humans suffering from social impairments.

To explore these mechanisms, we designed a social decision-making setup for pairs of macaques to simulate various social scenarios. The monkeys faced each other across a transparent touchscreen and performed a shape-cue decision making task. We aimed to investigate how monkeys behave in different situations, how their social hierarchies and familiarity may modulate their choices, and whether macaques consider the well-being of others when making choices leading to negative or positive outcomes for others. The setup allowed us to vary the probability of the other partner receiving a reward while keeping the probability of one’s own reward constant, testing the actor’s reaction to inequitable outcomes. Additionally, we incorporated a condition in which subjects had to decide to help others without receiving any reward. The setup was complemented by the use of cameras to monitor their behaviours and pair interactions, eye trackers to record social cues, and fNIRS to capture neural correlates. Dominance and social closeness were examined as contextual factors. This work establishes a foundation for understanding primate social decision-making, and provides valuable insights into the underlying mechanisms of social decision-making deficits in human disorders, potentially informing the development of future therapeutic interventions.

## 2. Results

We designed two tasks in which each participant in a dyad with known social rank performed either an object-categorisation or object-manipulation task where choices led to socially contingent reward outcomes. The contingent rules resulted in the following reward outcomes: reward–to– self, reward–to–both, reward–to–other, or reward–to–none. Subjects were free to chose among the actions presented in each specific social context.

For object categorisation (**Fig. 1b**), object shapes were associated with the reward outcomes — ★ = self | ⬣ = both | ▴ = other | ● = none. The positions, rotations and colours of the shapes were randomised on each trial so that subjects had to deliberately discount all other visual features when categorising the objects and selecting their preferred outcome. For object manipulation, a ball which obeyed physically simulated laws of gravity could be “flung” towards a correct or incorrect target, and the social outcome depended on the session context which was randomised from day to day (**Fig. 1c**). We measured the number of correct trials compared to the total number of attempted trials based on the social context.

**Figure 1.**
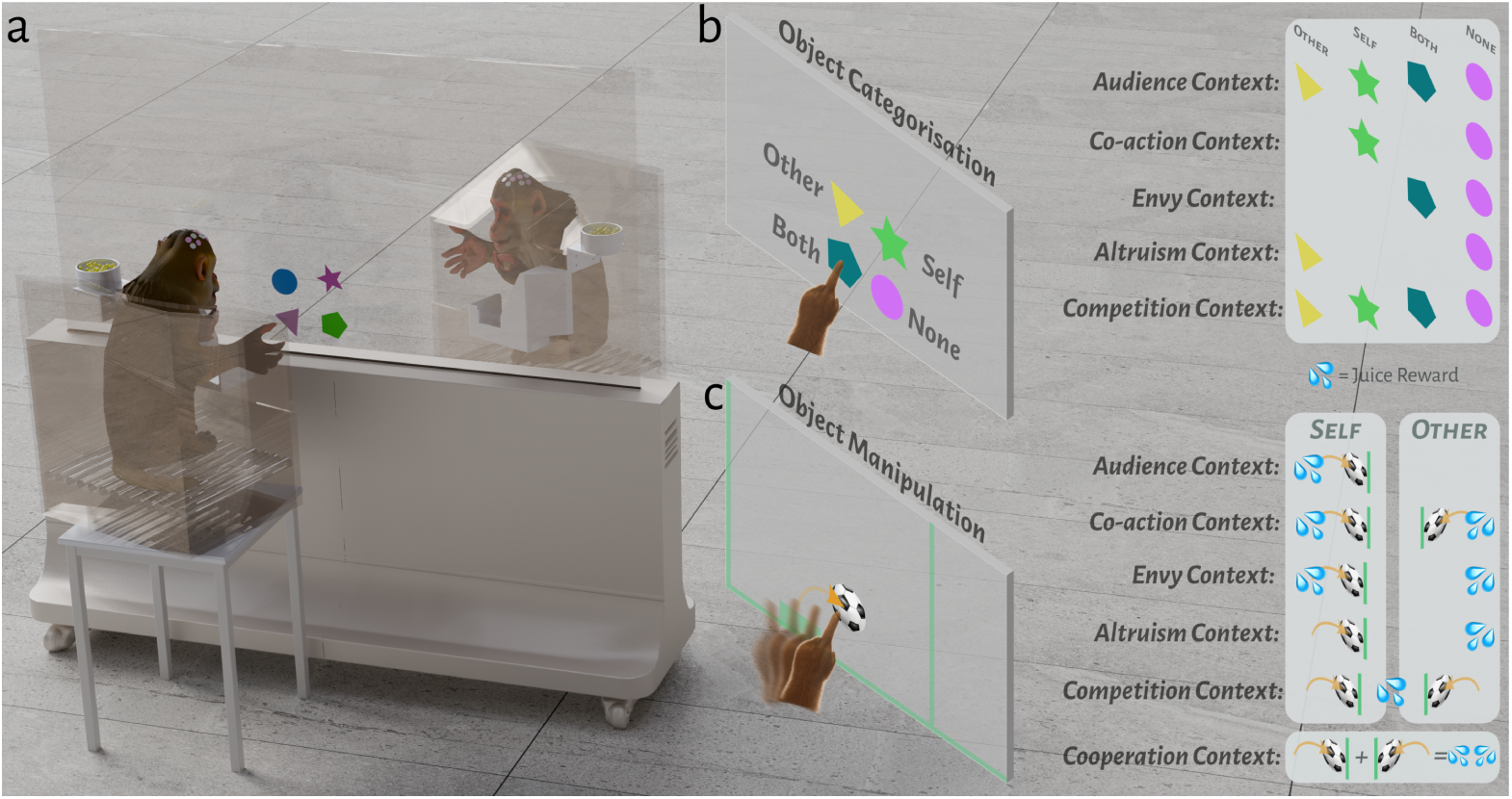
Social Dyad Decision Making Task. (**a**) experimental setup with two partners of a known social rank interacting via touch of a shared visual display. Dual eye-tracking and fNIRS recordings provided behavioural and neurophysiological indicators of task-related activity. Object categorisation task: shape symbolised reward contingencies to self and/or other; colours, angles and positions were randomised. By presenting different shapes within a block we could change the social context between audience, co-action, envy, altruism and competitive contexts. **(b)** Object manipulation task: virtual balls obeying newtonian physical laws were flung towards a divider. Rewards were given when the ball hit the divider, and by presenting shared or individual balls along with reward contingencies different social contexts could be created. This task enabled use to explore cooperative task where the ball had to be flung by partner A to partner B and back from partner B before both subjects could receive a reward.

### 2.1. Behavioural Results

#### 2.1.1. Decision Making Performance in Control and Social Conditions

The percentage of correct responses was not normally distributed (Shapiro-Wilk test, p < 0.05), but the variances were homogeneous (Mauchly’s test, p = 0.152, χ²(20) = 27.185). To compare the percentage of correct responses across conditions, we conducted a Friedman test, a non-parametric alternative to a repeated measure ANOVA test across conditions. The Friedman test revealed a significant difference between conditions (p = 4.852 × 10⁻⁸, χ²(6) = 44.92, Kendall’s W = 0.576), and a Bayesian analysis supports this effect (BF₁₀ = 3.328 × 10¹³).

To further investigate the differences between conditions, we performed Conover’s post-hoc comparison tests. The results showed that the macaques exhibited enhanced performance (p = 0.006, BF₁₀ = 16.07) in the presence of an audience conspecific compared to the control condition (**Fig. 2**). Conversely, the percentage of correct responses in the altruism and competition conditions was significantly lower than in the control, audience, coaction, envy, and cooperation conditions (p < 0.001, BF₁₀ > 10 for all comparisons, providing strong evidence).

**Figure 2.**
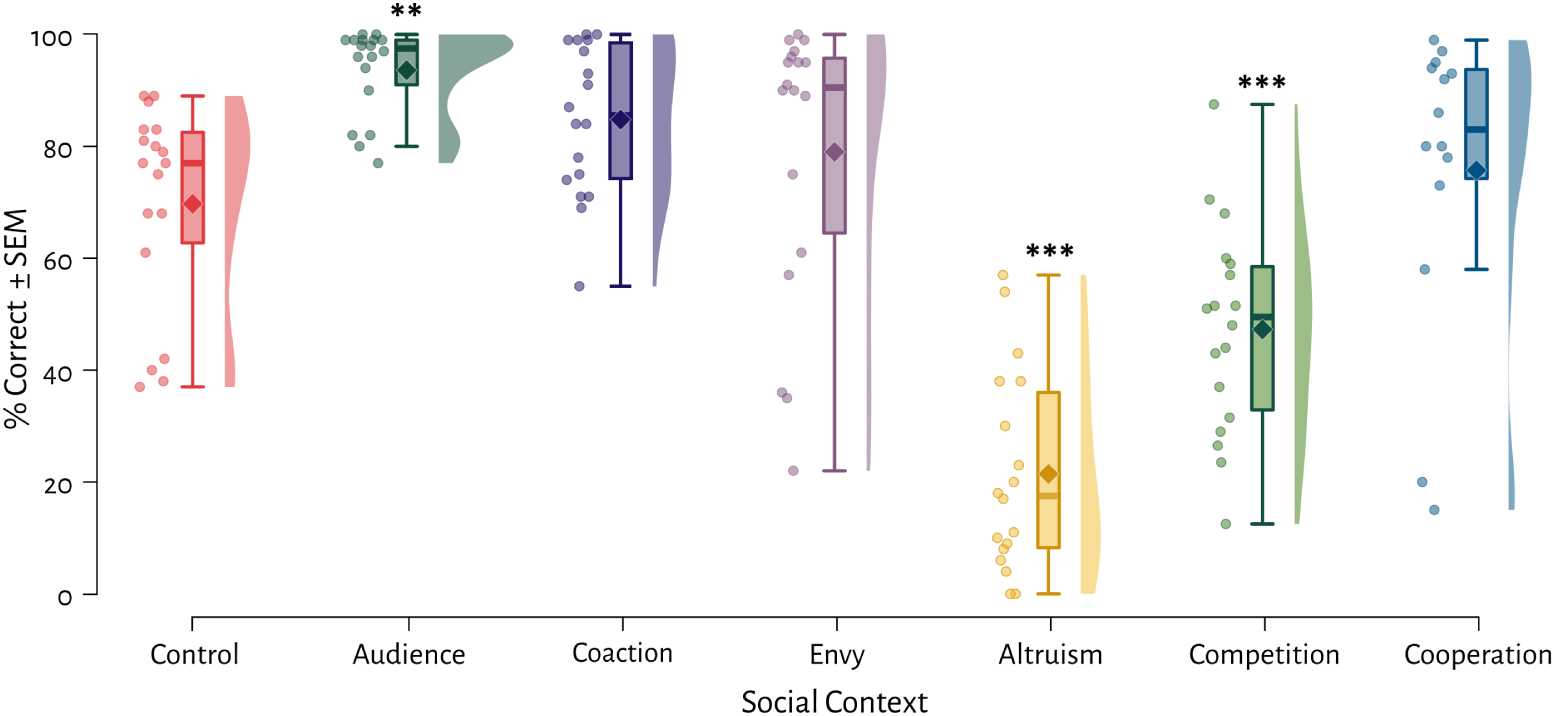
Percentage of correct responses in control and social conditions. The audience condition resulted in significantly higher correct response rates compared to the control condition, while the altruism and competition conditions exhibited significantly lower performance compared to other conditions. Comparisons shown versus control condition only; refer to text for other significant differences. **p<0.01, ***p<0.001 vs control.

We also examined the effect of dominance on performance (**Fig. 3**, between-subjects effects). Dominance showed a significant simple main effect (p = 2.994 × 10⁻⁸, F(6) = 14.271) and between-subjects effects (p = 0.018, ω² = 0.217). Post-hoc comparisons (Bonferroni-Holm corrected to control Type I error across all post-hoc dominance comparisons, as Friedman’s test does not handle between-subjects factors, and skew < 1, and kurtosis < 2, despite partial violations of sphericity (Mauchly’s p = 0.045)) revealed significant dominance effects in the Envy condition (mean difference = −30.05, p = 0.028, Cohen’s d = −1.70), Altruism condition (mean difference = −26.98, p = 0.007, d = −1.53), and Cooperation condition (mean difference = −31.41, p = 0.038, d = −1.78), with subordinate individuals outperforming dominants.

**Figure 3.**
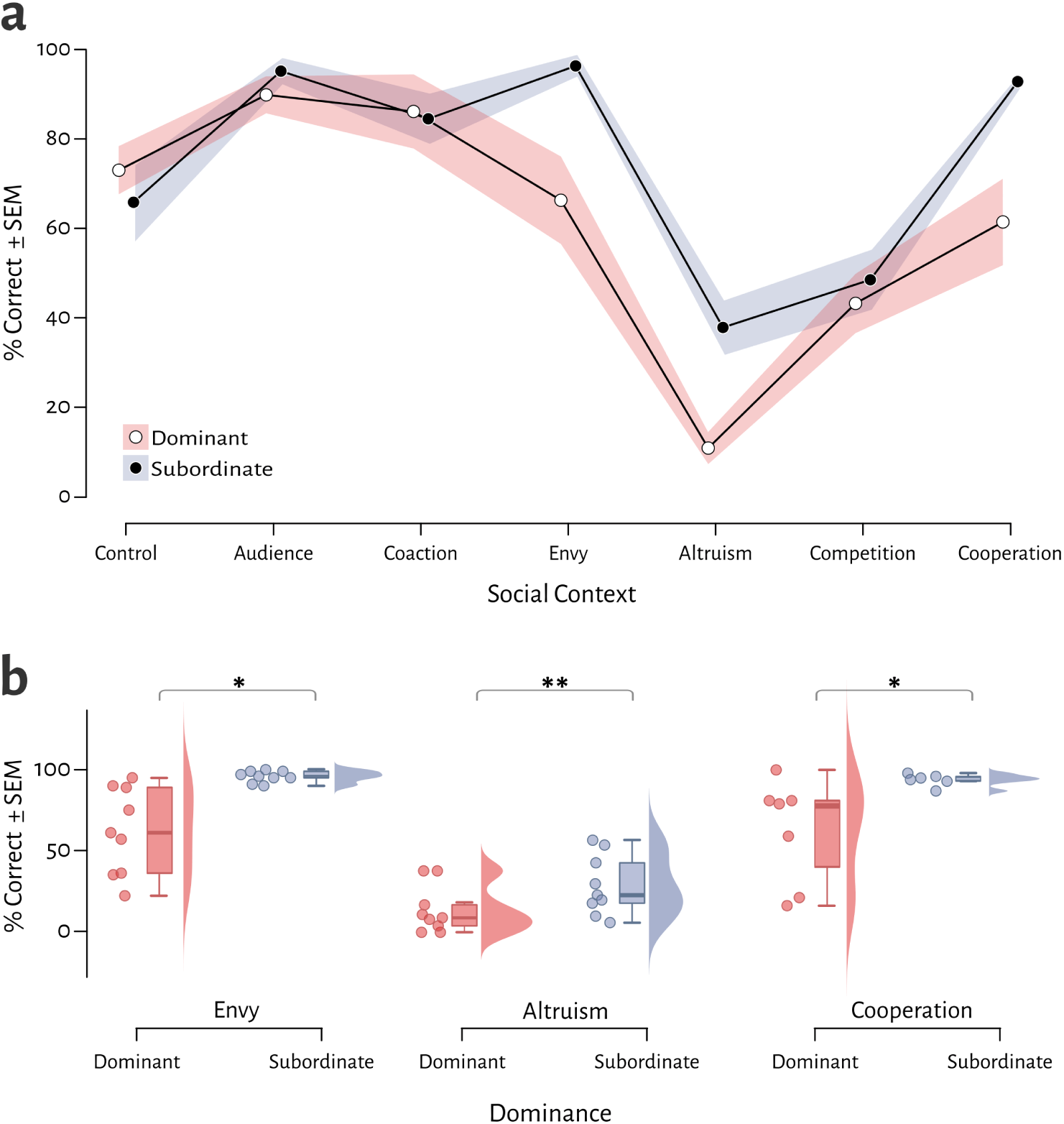
The effect of dominance on performance across conditions. (**a**) Overall response across social contexts. Note that in the control condition (where there is no conspecific, we used the overall rank to split the data) (**b**) Subordinate monkeys outperformed dominant monkeys in envy, altruism, and cooperation conditions. Gray bands represent ±SEM. *p<0.05, **p<0.01.

Due to non-normality (Shapiro-Wilk p < 0.05 in some conditions), we confirmed dominance effects with Mann-Whitney U tests (Bonferroni-Holm corrected, α = 0.007). Results aligned with parametric findings: for Envy (U = 3.5, p = 0.001, rank-biserial effect size = −0.914); for Altruism, a trend that was non-significant after correction (U = 18.0, p = 0.052, rank-biserial = −0.556); and for Cooperation (U = 6.0, p = 0.038, rank-biserial = −0.714).

This suggests that dominant monkeys performed worse when they completed the test correctly but both monkeys received a reward (Envy condition), when they had to decide to help their partner without any reward for themselves (altruism condition), or when they had to cooperate with a subordinate to jointly earn a reward (Cooperation condition).

#### 2.1.2. Reaction Times

The mean reaction time (RT), calculated as the average response duration for correct trials, violated assumptions of normality (Shapiro-Wilk p < 0.05) and sphericity (Mauchly’s test: χ²(14) = 27.32, p = 0.028). A Friedman test revealed significant differences across conditions (χ²(5) = 18.93, p = 0.002, Kendall’s W = 0.473), with Bayesian analysis corroborating this effect (BF₁₀ = 1445.492).

Post-hoc comparisons (Conover’s test) indicated that RTs were significantly faster in the competition condition compared to the control (p = 0.014, BF₁₀ = 2.17). The competition condition also elicited quicker responses than all other conditions: coaction (p = 2.741 × 10⁻⁵, BF₁₀ = 4830.03), envy (p = 0.002, BF₁₀ = 8.65), and altruism (p = 2.119 × 10⁻⁴, BF₁₀ = 5.22). Conversely, RTs were slowest in the coaction condition, which differed significantly from both the control (p = 0.032, BF₁₀ = 1.13) and audience conditions (p = 0.002, BF₁₀ = 784.38). The audience condition further showed faster RTs relative to the altruism condition (p = 0.014, BF₁₀ = 3.52).

In summary, the competition condition produced the shortest RTs, while coaction resulted in the longest, highlighting a robust effect of social context on response speed (**Fig. 4**).

**Figure 4.**
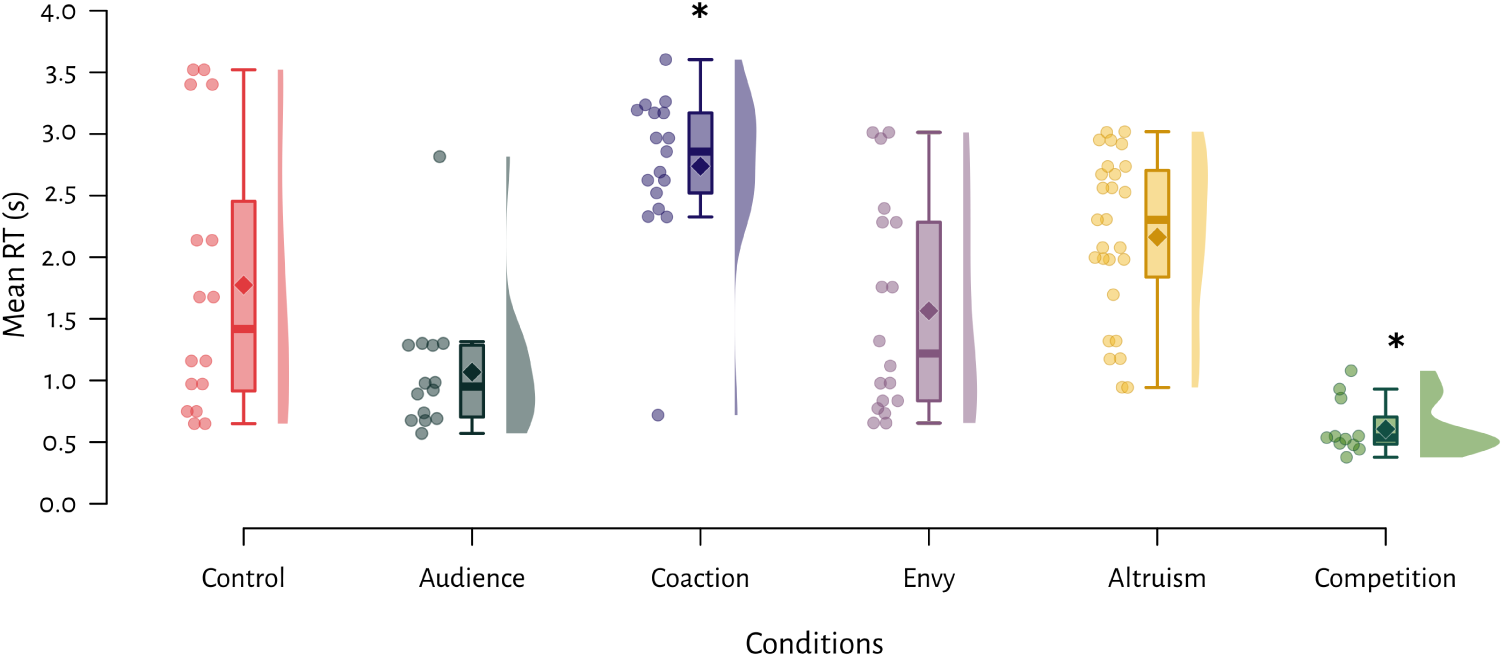
Mean reaction times for correct trials. Coaction significantly slower than control, and competition significantly faster than control. Comparisons shown versus control condition only. *p<0.05 vs control.

### 2.2. Social Gaze and Pupil Size Results

#### 2.2.1. Pupil Size

We measured pupil size as an index of arousal during task performance. The pupil size data met assumptions of normality (Shapiro-Wilk test, p > 0.05) and sphericity (Mauchly’s test: χ²(14) = 11.341, p = 0.723). Repeated-measures ANOVA revealed a significant main effect of condition on pupil size (F(5, 30) = 5.412, p = 1.929 × 10⁻⁵, ω² = 0.479), with strong Bayesian evidence supporting this effect (BF₁₀ = 4491.268).

Post-hoc Bonferroni-corrected comparisons demonstrated significantly larger pupil diameters in social conditions compared to control: audience (p = 0.048, Cohen’s d = −2.614, BF₁₀ = 13.777), coaction (p = 0.068, Cohen’s d = −2.299, BF₁₀ = 9.712), envy (p = 0.037, Cohen’s d = −2.038, BF₁₀ = 17.789), altruism (p = 0.009, Cohen’s d = −2.696, BF₁₀ = 60.223), and competition (p = 0.010, Cohen’s d = −3.157, BF₁₀ = 52.205). These robust effects indicate substantially heightened arousal when monkeys interacted with or observed conspecifics compared to solitary control trials (**Fig. 5a**).

**Figure 5.**
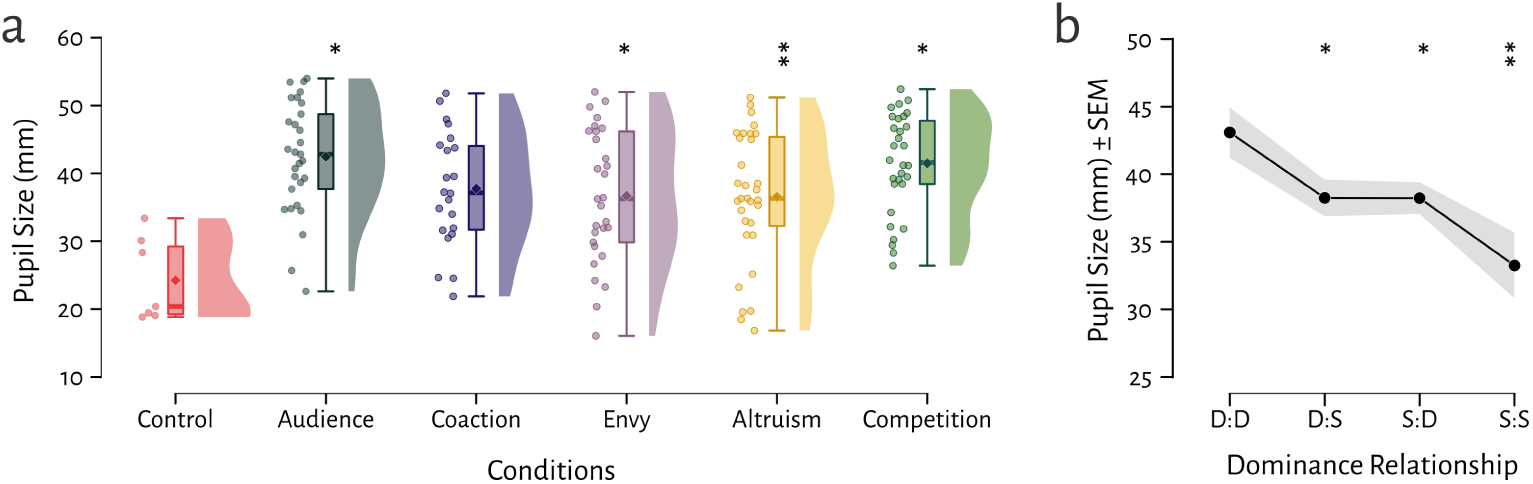
Pupil size changes across conditions. (**a**) All social conditions showed larger pupil diameters compared to control, indicating heightened arousal level. (**b**) Dominant-dominant pairs showed larger pupil diameters than other dominance combinations, indicating heightened arousal when two dominant monkeys performed together. Gray bands represent ±SEM. *p<0.05, **p<0.01.

#### 2.2.2. Pupil Size and Dominance

We also examined the effect of dominance on pupil size when two monkeys from different pairs performed the task together. The pupil size data were not normally distributed (Shapiro-Wilk test, p < 0.05), but the variances were homogeneous (Mauchly’s test, p = 0.279, χ²(5) = 6.31). The Friedman test revealed a significant difference in dominance (p = 0.007, χ²(3) = 12.22, Kendall’s W = 0.255), with Bayesian analysis corroborating this effect (BF₁₀ = 6.048). Conover’s post-hoc tests showed that pupil size was significantly larger when two dominant monkeys performed together than when two subordinate monkeys performed together (p = 3.830 × 10⁻⁴, BF₁₀ = 20.31), when a subordinate subject worked as an actor with a dominant monkey as a partner (p = 0.037, BF₁₀ = 0.34), or in the reverse scenario (p = 0.012, BF₁₀ = 2.40). This suggests that dominant monkeys experience greater arousal when competing with another dominant monkey (Rosvold *et al*., 1954) (**Fig. 5b**).

#### 2.2.3. ROI Time Percent

#### 2.2.4. iRecHS2 ROI looking time across conditions

We additionally evaluated the amount of time that monkeys spent looking at their partner’s face region on the other side (ROI time data). The ROI time data were not normally distributed (Shapiro-Wilk test, p < 0.05), but the sphericity assumption was met (Mauchly’s test, p = 0.782, χ²(14) = 10.184). The Friedman test revealed a significant effect of condition (p = 0.012, χ²(5) = 14.714, Kendall’s W = 0.368), with Bayesian analysis corroborating this effect (BF₁₀ = 4.655).

Post-hoc Conover tests showed that ROI time was significantly greater in all conditions than in the control condition (envy: p = 0.003, BF₁₀ = 14.79; competition: p = 0.014, BF₁₀ = 8.02; coaction: p = 0.046, BF₁₀ = 2.26; audience: p = 0.035, BF₁₀ = 5.33; altruism: p = 0.004, BF₁₀ = 10.00). This suggests that monkeys were interested in their partner’s behaviour when engaged in social tasks (**Fig. 6a**).

**Figure 6.**
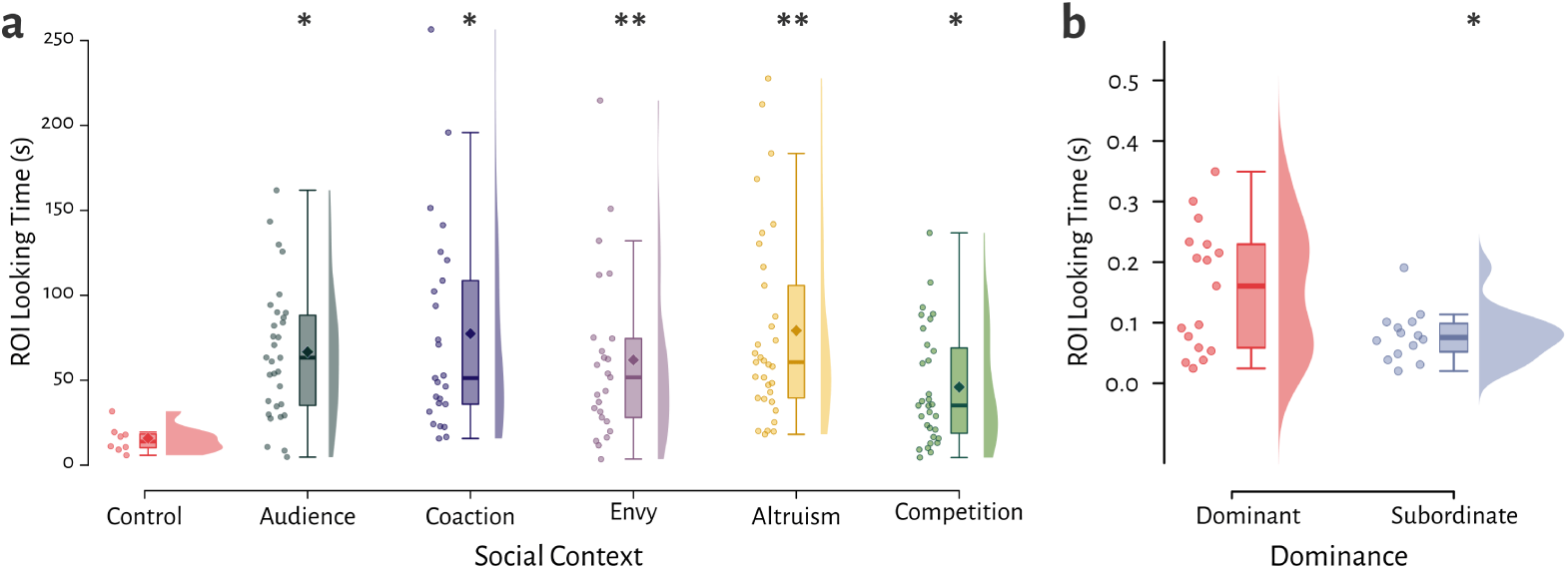
ROI looking across conditions. (**a**) All social conditions showed significantly greater ROI looking time than control, suggesting that monkeys are interested in their partner’s behaviour when they are engaged in social tasks. *p<0.05, **p<0.01 vs control. (**b**) Dominant monkeys exhibited significantly higher face-looking time than subordinates. *p<0.05.

We additionally evaluated dominance in ROI time percentage in each condition. This was calculated as the proportion of time the subject spent looking at the ROI region compared to looking outside of the ROI region. The data were normally distributed (Shapiro-Wilk test, p > 0.05), but the variances were not homogeneous (Levene’s test, p = 1.38 × 10⁻⁴). We found a significant difference between dominant and submissive monkeys in the competition condition (Welch test, p = 0.011, BF₁₀ = 3.78). Dominant monkeys spent more time looking at their partner’s face in the competition condition than submissive monkeys (**Fig. 6b**). This may be because dominant monkeys had greater expectations to gain from winning competitions, because they were more motivated to avoid losing, and/or the fact that subordinate monkeys tend to avoid looking at other monkeys’ face/eye regions to minimise interaction.

We also observed that dominant monkeys displayed aggressive behaviour when a submissive monkey touched the visual cue first and received the reward. This is consistent with the point that dominant monkeys can use aggression to maintain their social status or discourage other monkeys from performing the task faster!

#### 2.2.5. Saccade Frequency to Partner’s Face (ROI) Across Task Phase

We evaluated the average number of saccades per trial during pre-stimulus, response time, and post-stimulus periods, quantifying how frequently monkeys looked at their partner’s face region (ROI) across conditions. The data violated normality assumptions (Shapiro-Wilk test, p < 0.05), prompting use of Friedman tests which revealed significant condition differences (χ²(14) = 33.118, p = 0.003, Kendall’s W = 0.215), with decisive Bayesian evidence (BF₁₀ = 3415.347).

Our analysis (**Fig. 7a**) examined both between-condition differences (comparing the same interval across different conditions) and within-condition differences (comparing different intervals within the same condition). Between-condition comparisons showed significant pre-stimulus differences: audience versus competition (p = 0.020, r_r_b = −0.788, BF₁₀ = 1.690), envy versus competition (p = 3.791×10⁻⁴, r_r_b = −0.848, BF₁₀ = 3.292), and altruism versus competition (p = 0.029, r_r_b = −0.970, BF₁₀ = 6.494).

**Figure 7.**
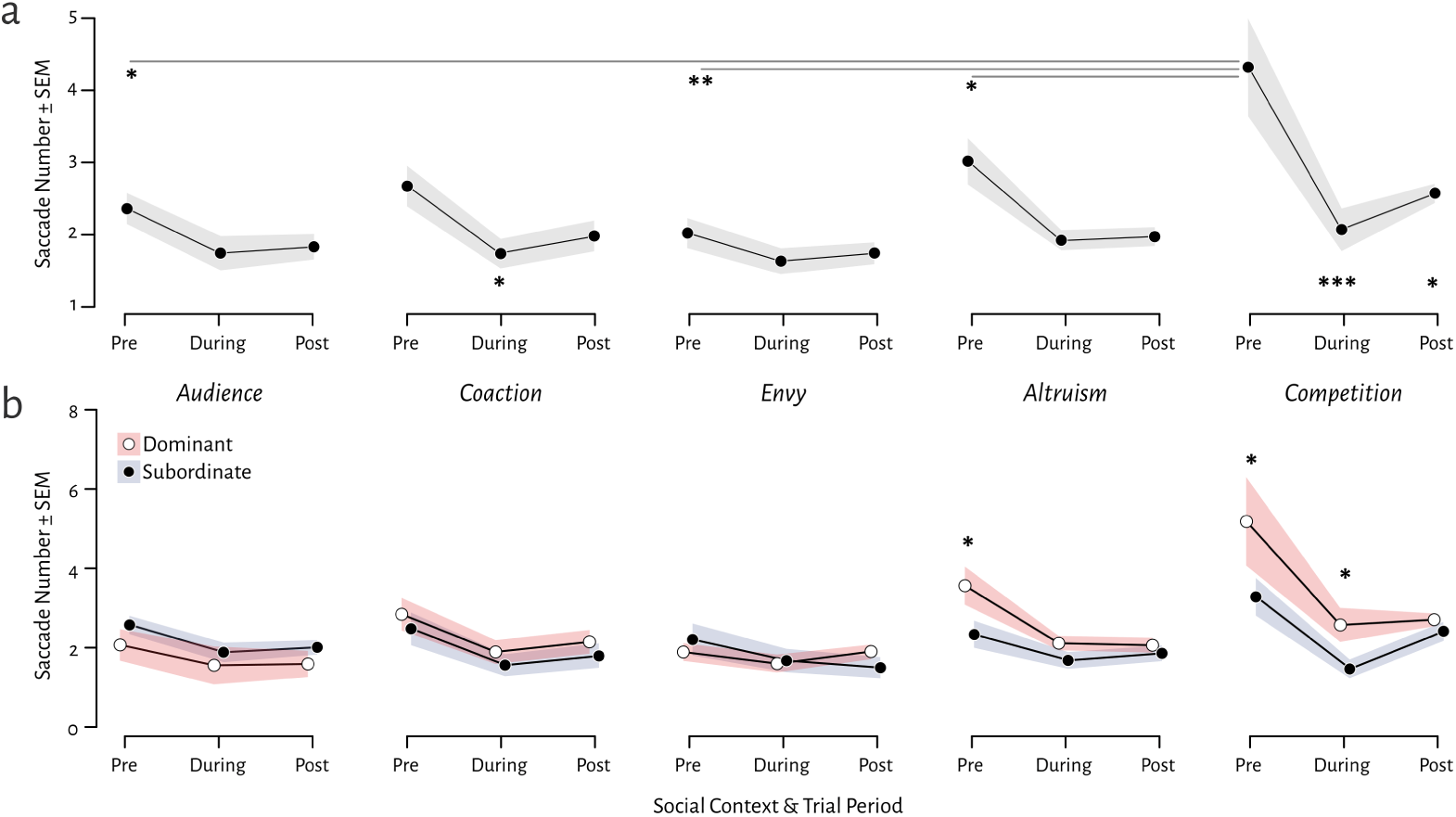
Saccade frequency to partner’s face (ROI) across task phases and conditions. (**a**) Pre-decision saccades were more frequent than during/post-response in coaction and competition conditions, indicating heightened social monitoring before choices. Pre-decision saccades between conditions were larger in competition compare to audience, envy, and altruism. (**b**) Dominant monkeys showed increased looking during pre-stimulus phases in competitive and altruistic conditions compared to subordinates. *p<0.05, **p<0.01.

Within-condition comparisons revealed significant variations across time periods in specific conditions. For coaction, pre-stimulus fixation was significantly higher than response-period fixation (p = 0.042, r_r_b = 1, BF₁₀ = 51.376). In the competition condition, pre-stimulus fixation was higher than both response-period (p = 3.158×10⁻⁴, r_r_b = 0.939, BF₁₀ = 6.822) and post-stimulus fixation (p = 0.047, r_r_b = 1, BF₁₀ = 2.932). This trend indicates that monkeys typically made more saccades to the ROI (reflecting greater social monitoring) prior to their decisions.

We also examined dominance effects on performance (as between-subjects effects). Dominance showed a significant main effect (F(14) = 5.973, p = 5.209×10⁻¹⁰). Due to non-normality (Shapiro-Wilk p < 0.05 in some conditions) and Friedman test limitations for between-subjects comparisons, we analysed dominance effects using Mann-Whitney U tests. Results revealed significant differences between dominant and subordinate monkeys in: altruism pre-stimulus periods (p = 0.020, rank-biserial = 222.0, BF₁₀ = 2.023); competition pre-stimulus (p = 0.040, rank-biserial = 64.00, BF₁₀ = 1.118); and competition during performance (p = 0.022, rank-biserial = 73.00, BF₁₀ = 3.06).

The trends suggest dominant monkeys monitored their partners more frequently, particularly during pre-decision periods and in competitive contexts (competition condition), as well as when deciding to provide help without reciprocal reward (altruism condition) (**Fig. 7b**).

#### 2.2.6. Pupil Core Total Fixation Time to ROI (Partner’s Face Region)

We analysed the total time monkeys spent fixating on their partner’s face region (ROI) across experimental conditions. The ROI fixation time data violated normality assumptions (Shapiro-Wilk test, p < 0.05). PupilCore measurements showed similar trends to iRecHS2 data in fixation patterns, with Friedman tests revealing statistically significant differences in face-looking time across conditions (χ²(6) = 14.743, p = 0.022, Kendall’s W = 0.491). Bayesian repeated measures ANOVA further supported these condition differences (BF₁₀ = 4.357).

Post-hoc analyses demonstrated that control condition fixation times differed significantly from audience (p = 0.001, r_r_b = −1.000, BF₁₀ = 935.037), coaction (p = 0.017, r_r_b = −1.000, BF₁₀ = 20.050), envy (p = 0.002, r_r_b = −1.000, BF₁₀ = 5.008), and competition (p = 0.001, r_r_b = −1.000, BF₁₀ = 16.030) conditions, with a marginal difference for cooperation (p = 0.055, r_r_b = −1.000, BF₁₀ = 93.684). The altruism condition also showed significant differences compared to audience (p = 0.025, r_r_b = −1.000, BF₁₀ = 1.007), envy (p = 0.037, r_r_b = −0.733, BF₁₀ = 0.839), and competition (p = 0.025, r_r_b = −0.867, BF₁₀ = 0.481) conditions (**Fig. 8**).

**Figure 8.**
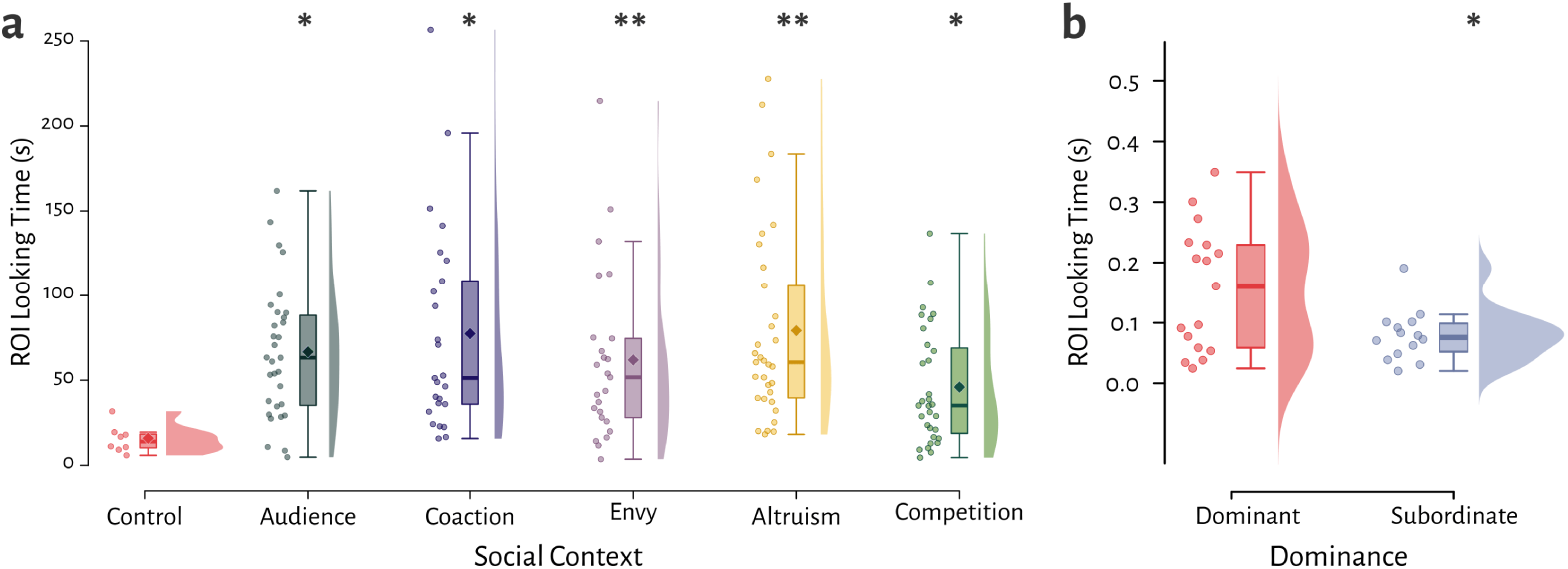
Total fixation time to partner’s face (ROI). Almost all social conditions showed significantly longer looking times than control, indicating heightened interest in partner’s behaviour during social tasks. Comparisons shown versus control condition only. *p<0.05,**p<0.01 vs control.

#### 2.2.7. Blink Rate Modulation Across Social Contexts

Spontaneous eye-blinks have been proposed to function as a social signal in primates, with studies suggesting their role in communication parallels that of grooming in complex social interactions (Tada *et al*., 2013). Our blink frequency data violated normality assumptions (Shapiro-Wilk p < 0.05). A Friedman test revealed significant differences across conditions (χ²(6) = 22.071, p = 0.001, Kendall’s W = 0.613), with Bayesian analysis providing decisive evidence for these effects (BF₁₀ = 4368.395).

Post-hoc comparisons demonstrated that blink rates in the coaction (p = 0.002, BF₁₀ = 2.083) and competition (p = 3.064×10⁻⁵, BF₁₀ = 6.590) conditions were significantly lower than in the control condition.

The audience condition also showed significantly different blink frequencies compared to both coaction (p = 0.010, BF₁₀ = 1.613) and competition (p = 1.598×10⁻⁴, BF₁₀ = 28.371) conditions. Notably, coaction elicited significantly different blink patterns from envy (p = 4.725×10⁻⁴, BF₁₀ = 5.450), altruism (p = 0.025, BF₁₀ = 5.514), and cooperation (p = 0.006, BF₁₀ = 14.187) conditions. The most robust differences emerged in competitive contexts, where blink rates in the competition condition differed sharply from those in the envy (p = 5.820×10⁻⁶, BF₁₀ = 205.353), altruism (p = 4.725×10⁻⁴, BF₁₀ = 164.510), and cooperation (p = 9.239×10⁻⁵, BF₁₀ = 2661.885) conditions (**Fig. 9**).

**Figure 9.**
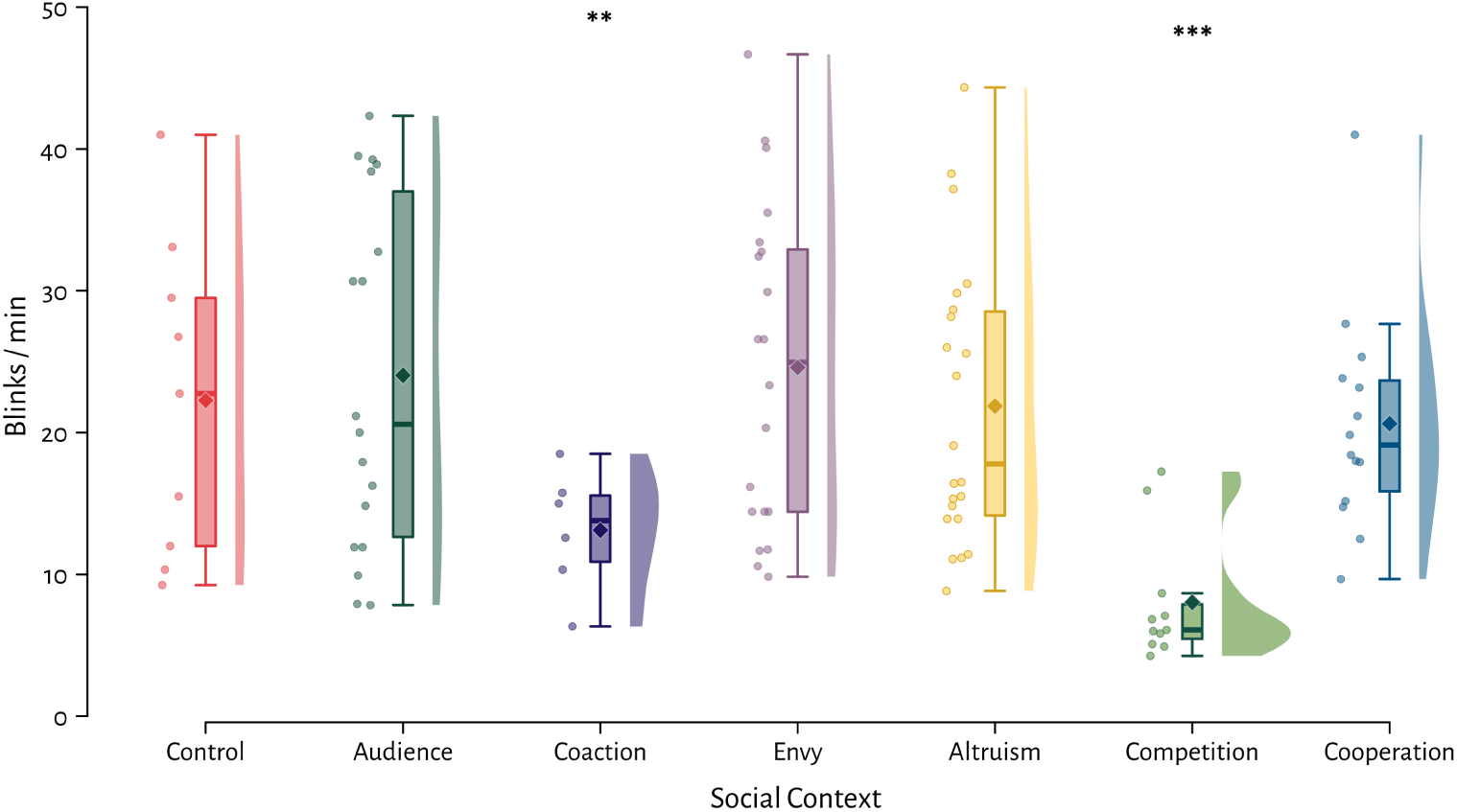
Blink rates during task performance. Coaction and competition conditions showed strongest modulation versus control during simultaneous partner monitoring. Comparisons shown versus control condition only; refer to text for other significant differences. **p<0.01, ***p<0.001 vs control.

The most behaviourally significant results occurred during simultaneous task performance, when monkeys actively monitored their partner’s actions while engaged in the task.

### 2.3. fNIRS Haemodynamic Responses Across Social Conditions

#### 2.3.1. Overall Haemodynamic Responses Across Fronto-temporal Areas

**HbO Analysis**: The fNIRS HbO data violated assumptions of normality (Shapiro-Wilk p < 0.05) and sphericity (Mauchly’s test: χ²(20) = 35.311, p = 0.019). Friedman tests revealed significant between-condition differences (χ²(6) = 21.646, p = 0.019, Kendall’s W = 0.023), with Bayesian support (BF₁₀ = 36.868). Post-hoc comparisons demonstrated that the control condition differed significantly from envy (p = 0.007, r = 0.257, BF₁₀ = 2.585), altruism (p = 0.003, r = 0.353, BF₁₀ = 339.656), and competition (p = 0.013, r = 0.293, BF₁₀ = 31.499) (fig. 17).

The audience condition showed distinct responses compared to envy (p = 0.020, r = 0.176, BF₁₀ = 0.192), altruism (p = 0.011, r = 0.252, BF₁₀ = 2.473), and competition (p = 0.036, r = 0.179, BF₁₀ = 1.346). Coaction responses differed from envy (p = 0.003, r = 0.309, BF₁₀ = 1.700), altruism (p = 0.001, r = 0.353, BF₁₀ = 152.697), and competition (p = 0.006, r = 0.274, BF₁₀ = 9.573).

**HbR Analysis**: HbR data similarly violated normality (p < 0.05) and sphericity (Mauchly’s test: χ²(20) = 37.145, p = 0.011). Significant between-condition differences were observed (χ²(6) = 22.358, p = 0.001, Kendall’s W = 0.023), though Bayesian evidence was weaker (BF₁₀ = 2.836). Control condition responses differed from coaction (p = 0.013, r = −0.169, BF₁₀ = 0.220), envy (p = 0.029, r = −0.267, BF₁₀ = 2.514), altruism (p = 0.010, r = −0.270, BF₁₀ = 3.779), and competition (p = 4.521×10⁻⁵, r = −0.387, BF₁₀ = 516.459) (fig. 17).

Additional differences emerged between audience and competition (p = 0.008, r = −0.286, BF₁₀ = 2.285), as well as between cooperation and altruism (p = 0.042, r = −0.191, BF₁₀ = 0.635) and competition (p = 4.042×10⁻⁴, r = −0.309, BF₁₀ = 32.129).

**HbT Analysis**: HbT data were non-normal (Shapiro-Wilk test, p < 0.05) with heterogeneous variances (Mauchly’s test: χ²(20) = 40.441, p = 0.004). Friedman tests showed no significant between-condition differences (χ²(6) = 7.931, p = 0.243, Kendall’s W = 0.008), supported by Bayesian evidence for the null (BF₁₀ = 0.015) (**Fig. 10**). The absence of significant differences in total haemoglobin (HbT) across these social conditions is also consistent with fNIRS expectations, as HbT changes are often smaller and less responsive to task execution compared to the differential changes in HbO and HbR.

**Figure 10.**
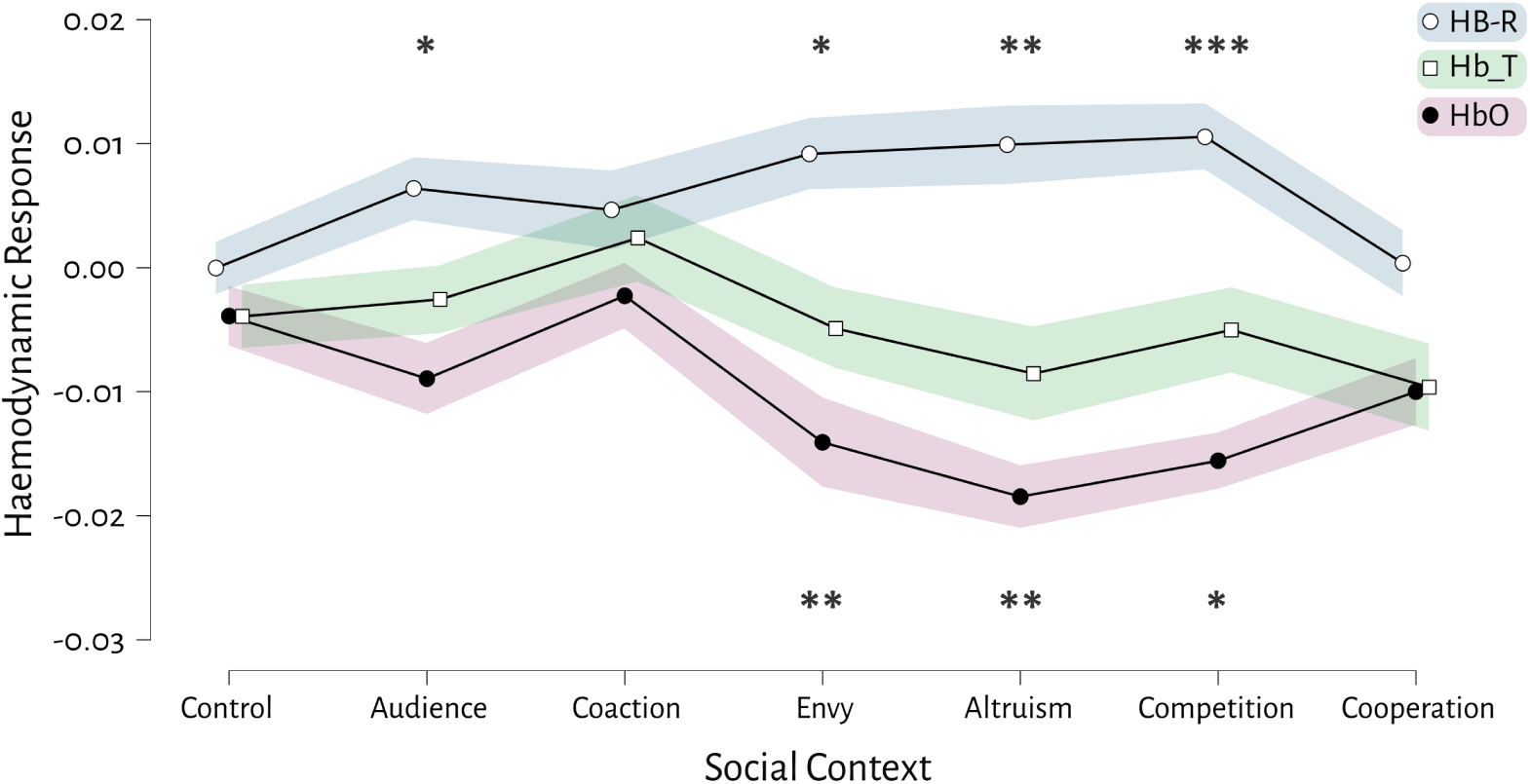
fNIRS haemodynamic responses versus control. HbO: Social conditions (coaction, envy, altruism, and competition) > control. HbR: Envy, altruism, competition < control HbT: No condition differences detected. Comparisons shown versus control only; see text for other contrasts. *p<0.05, **p<0.01, ***p<0.001.

#### 2.3.2. General Linear Model (GLM) Analysis of Evoked fNIRS Data

Due to the monkeys’ limited ability to sustain prolonged task engagement, we implemented a rapid event-related design (3-second task duration, 4-second inter-trial interval) instead of a traditional block design (**Supp. Fig. 1**), which typically requires longer intervals (7–8 s task, 10–15 s rest) for optimal haemodynamic response function (HRF) estimation (Huppert *et al*., 2009). To analyse these evoked responses, we employed a GLM, a validated method for fNIRS rapid designs (Plichta *et al*., 2007; Tak & Ye, 2014). The GLM effectively modelled the HRF while compensating for overlapping haemodynamic responses, controlled for physiological noise and motion artefacts (Barker *et al*., 2013), and isolated task-evoked neural activity from confounding factors to enable robust condition comparisons.

We modelled the haemodynamic response for each condition using the GLM with separate regressors for each experimental condition (Audience, Competition, Control, Envy, Coaction, altruism, Cooperation). The design matrix incorporated task regressors convolved with a canonical hemodynamic response function (HRF) along with nuisance regressors accounting for physiological noise and motion artefacts. Each condition-specific regressor was time-locked to its respective onset and convolved with a canonical HRF (3-second duration). Nuisance regressors included six head motion parameters (three translations + three rotations), bandpass-filtered physiological noise (0.01-0.2 Hz), and a first-order polynomial drift term.

Analysis of beta weights revealed systematic variation in task-evoked responses (Beta₁) across conditions, with Beta₂ accounting for nuisance variance. All analyses were implemented in the nirSpark fNIRS Analysis Toolkit (v1.8.8). Shapiro-Wilk tests confirmed non-normality (p < 0.05 across multiple conditions), with Mauchly’s test indicating variance heterogeneity (χ²(20) = 161.42, p = 4.288 × 10⁻²⁴), justifying our use of non-parametric approaches.

Friedman’s test showed highly significant between-condition differences in Beta₁ values (χ²(6) = 59.93, p = 4.654 × 10⁻¹¹, W = 0.104), with Bayesian signed-rank tests providing decisive evidence for these effects (BF₁₀ = 64,878.539). Post-hoc analyses revealed that the control condition showed significantly lower Beta₁ values compared to audience (p = 0.011, r = −0.329, BF₁₀ = 0.321), altruism (p = 3.767 × 10⁻⁶, r = −0.510, BF₁₀ = 2,004.489), competition (p = 5.304 × 10⁻⁵, r = −0.319, BF₁₀ = 1.213), and cooperation (p = 0.030, r = −0.218, BF₁₀ = 0.264) conditions, with altruism showing the highest Beta₁ values among these comparisons. The audience condition exhibited intermediate Beta₁ values, being significantly higher than coaction (p = 5.304 × 10⁻⁵, r = −0.522, BF₁₀ = 25.112) and envy (p = 0.003, r = −0.334, BF₁₀ = 2.529) but lower than altruism (p = 0.036, r = −0.243, BF₁₀ = 1.005).

Notably, coaction showed among the lowest Beta₁ values, being significantly different from conditions with higher responses including altruism (p = 1.226 × 10⁻⁹, r = −0.605, BF₁₀ = 307,317.256), competition (p = 3.672 × 10⁻⁸, r = −0.495, BF₁₀ = 225.124), and cooperation (p = 2.491 × 10⁻⁴, r = −0.407, BF₁₀ = 92.708). Similarly, the envy condition demonstrated relatively low Beta₁ values compared to altruism (p = 4.848 × 10⁻⁷, r = −0.497, BF₁₀ = 154.077), competition (p = 8.466 × 10⁻⁶, r = −0.385, BF₁₀ = 6.066), and cooperation (p = 0.010, r = −0.329, BF₁₀ = 1.712). The cooperation condition showed intermediate Beta₁ values, being significantly higher than control but lower than altruism (p = 0.013, r = −0.432, BF₁₀ = 20.865) and marginally higher than competition (p = 0.058, r = −0.231, BF₁₀ = 0.305).

## 3. Discussion

We designed a naturalistic social decision-making paradigm in which macaque pairs interacted face-to-face across a transparent touchscreen, completing tasks under varying social contexts while behavioural, ocular, and neuroimaging data were simultaneously collected. Our results demonstrate that macaques’ decision-making is dynamically modulated by social context. Subjects exhibited significantly enhanced cognitive performance in the audience condition compared to isolated control trials, suggesting that the mere presence of a conspecific sharpens attention— consistent with prior findings in humans (Ganesh *et al*., 2014) and monkeys (Chang *et al*., 2015; Reynaud *et al*., 2015). Conversely, performance declined sharply in the altruism condition (where no reward was received) and in the competition condition, where aggressive behaviours—particularly in dominant individuals—led to missed trials. Dominant individuals underperformed in envy, altruism, and cooperation conditions, where immediate rewards were absent or shared. These behavioural patterns is align with prior work showing that macaque decision-making integrates social evaluation (Deaner *et al*., 2005; Seyfarth & Cheney, 2014).

Physiological measurements revealed increased arousal across all social conditions, with dominance relationships further modulating pupillary responses and gaze patterns. Neural data confirmed activation in cortical regions associated with social decision-making, particularly in envy, altruism, and competition conditions, reinforcing their crucial role in these processes. Together, these findings illustrate how macaques integrate social variables with reward contingencies, with hierarchical relationships shaping both cognitive and physiological responses.

### 3.1. The Role of Vicarious Affect and Pro-sociality in Nonhuman Primates

The extent to which nonhuman primates experience vicariously induced affective states during social decision-making remains debated (Panksepp & Panksepp, 2013; Tan & Hare, 2013; Noritake *et al*., 2018). While some studies suggest primates exhibit greater prosocial behaviour (Cronin, 2012; Tan & Hare, 2013; Emigh *et al*., 2020), including activation of reward circuits when observing conspecifics receive rewards (Chang *et al*., 2011), our findings indicate that long-tailed macaques were reluctant to help conspecifics without personal reward. This aligns with studies showing that rhesus macaques avoid prosocial choices when they incur costs (Sallet *et al*., 2021), and that orangoutangs do not naturally share food even when it imposes no cost (Kim *et al*., 2015). Similarly, in group feeding settings, dominant rhesus macaques punish subordinates for approaching food, further illustrating the scarcity of altruism in reward-based contexts (Chancellor & Isbell, 2008).

These discrepancies may arise from differences in experimental design, species-specific traits, living conditions, or individual variation. For instance, prosocial behaviour in male mice is rare and limited to a small subset of individuals (Esteve-Agraz *et al*., 2025), suggesting that such tendencies even may not seen in all individuals of a species and are only observed in a small proportion of animals.

### 3.2. Familiarity and Dominance in Social Decision-Making

We also examined whether familiarity (cage-mate vs. non-cage-mate) influenced monkeys’ choices in our study. While several studies report partner selectivity in primates and rodents, with increased prosocial behaviour toward familiar individuals (Chang *et al*., 2011; Massen *et al*., 2011; Ben-Ami Bartal *et al*., 2014; Garcia *et al*., 2023), we observed no significant differences in our study. This null effect may be attributed to our subjects being long-term group-housed, having lived together for their first two years of life, which likely resulted in all pairs being highly familiar with one another. Previous research has demonstrated that familiarity enhances social interactions in both humans and monkeys (Preston & de Waal, 2002; Garcia *et al*., 2023), and that actors tend to favour both familiar and subordinate recipients (Batson & Powell, 2003; Chang *et al*., 2011). However, our results align with findings in rats where social hierarchy, but not familiarity, modulated prosocial choices (Gachomba *et al*., 2022).

Regarding dominance effects, our results revealed that subordinate monkeys outperformed dominant individuals in envy, altruism, and competition conditions. This finding contrasts with studies reporting greater pro-sociality in dominant individuals (Massen *et al*., 2010; Chang *et al*., 2011). Several factors may explain this discrepancy. First, methodological limitations in prior work should be considered; for instance, Chang et al. (2011) tested only two middle-ranked male macaques, and Ballesta et al. (2015) found that benevolent tendencies were exhibited exclusively by the single dominant monkey (M1) in their small sample study.

Second, contextual and species-specific factors likely play important roles. Dominant macaques may resist prosocial acts when such behaviours incur costs, such as reward sacrifice (Massen *et al*., 2010), whereas subordinates, lacking control over resources, may comply to avoid conflict (Chancellor & Isbell, 2008). Furthermore, differences between captive and natural hierarchies may be influential. In wild or colony settings, dominants may use prosocial behaviour to reinforce group cohesion (Massen *et al*., 2010; Seyfarth & Cheney, 2014), whereas in captive pairs like those in our study, social rank may remain stable without requiring such negotiation. Finally, species differences must be considered; while dominants demonstrate greater pro-sociality in one species (Gachomba *et al*., 2022), but not others (Chancellor & Isbell, 2008), with even different outcomes reported within the same species across studies. These findings collectively suggest that dominance-related prosocial behaviour is highly context-dependent, shaped by ecological pressures, experimental paradigms, and species-specific social strategies. Future studies would benefit from larger sample sizes and more naturalistic testing environments to further elucidate these complex relationships.

### 3.3. Behavioural Results

Our results demonstrate that face-area eye fixations were proportionally higher than non-face region fixations, aligning with Gilardeau et al. (2021) in showing that the partner’s face represents a highly salient visual feature. Regarding region-of-interest (ROI) analysis, we observed that direct head-fixation and mutual eye contact occurred infrequently, consistent with natural primate behaviour where such gaze patterns typically signal threat or aggression rather than typical social interaction. Notably, dominant monkeys exhibited more frequent and prolonged eye-region fixations compared to subordinates (Gilardeau *et al*., 2021), potentially reflecting their social monitoring role.

The interpretation of mutual gaze remains controversial in primate literature. While some studies characterise it as prosocial interaction or empathy indicator (Thomsen, 1974; Leonard *et al*., 2012; Ballesta & Duhamel, 2015; Gilardeau *et al*., 2021), our observations support the alternative view that eye contact primarily signals aggression in macaques (Fernandez *et al*., 2009). This was particularly evident during unrewarded trials (competition/altruism conditions), where gaze coincided with hostile behaviours. The amygdala activation reported during eye contact (Gilardeau *et al*., 2021) further supports this threat interpretation rather than prosocial signalling. Our data showed scarce mutual gaze, with frequent averted gaze when one monkey fixated on the eye region, consistent with Ballesta et al.‘s (2015) observations of natural avoidance patterns.

Our analysis revealed distinct temporal dynamics in gaze behaviour. We observed increased saccadic activity primarily during pre-stimulus periods and before task initiation, rather than during task execution or afterward. This pattern differs from Chang et al. (Chang *et al*., 2015), who reported that actors frequently looked at recipients after making decisions. In our paradigm, reward delivery occurred during inter-trial intervals (post-task), might leading subjects to shift attention to reward. This methodological difference likely accounts for the contrasting findings, as Chang et al. (Chang *et al*., 2015) employed separate pre-reward and post-reward time windows (split time windows to before and after task). Notably, we still observed empathic eye blinking patterns comparable to those reported by Ballesta et al. (Ballesta & Duhamel, 2015), suggesting this behaviour represents a conserved submissive signal across experimental paradigms.

These findings collectively highlight how gaze patterns in experimental settings reflect both natural communication systems and task-specific contingencies. The scarcity of direct eye contact supports ecological validity concerns raised by Cronin et al. (Cronin, 2016), while the observed fixation patterns nevertheless provide meaningful behavioural markers of social attention and hierarchy maintenance. Future studies should consider these gaze dynamics when designing social cognition paradigms for nonhuman primates.

### 3.4. Brain Areas Involved in Social Interactions

For fNIRS evaluation, we focused on dorsolateral prefrontal cortex (dlPFC), temporoparietal junction (TPJ), and superior temporal sulcus (STS), which are superficial cortical regions optimally accessible to optical neuroimaging. These areas constitute a core social decision-making network: the TPJ plays well-established roles in social cognition (Donaldson *et al*., 2015), particularly in processing others’ mental states (Wang *et al*., 2025), while STS integrates biological motion cues critical for social perception. The dlPFC, along with these regions, forms an integrated system for social reward evaluation, as demonstrated by primate studies showing dlPFC-V4 (Franch *et al*., 2024) and dmPFC-mSTS interactions during social information processing (Mahmoodi *et al*., 2024).

Deeper structures involved in social interactions (Tremblay *et al*., 2017) like the amygdala, anterior cingulate cortex (ACC), and ventral striatum undoubtedly contribute to social behaviour (Flechsenhar *et al*., 2022; Kim & Sul, 2023), with ACC neurons predicting conspecifics’ intentions during cooperation (Haroush & Williams, 2015) and evaluating social information with respect to others (envy and schadenfreude, Takahashi *et al*., 2009). The amygdala processes social cues (Wang *et al*., 2025). However, their depth limits fNIRS reliability. Our selection therefore balances anatomical accessibility with functional relevance, targeting regions where mirror neuron systems (Rizzolatti & Sinigaglia, 2010) and reward-prediction pathways (Burke *et al*., 2010) converge to transform social variables into decisions.

### 3.5. Haemodynamic Signatures

There are not a lot of fNIRS studies on primates, with most focusing on motor areas and often using anaesthetised animals (Yamada *et al*., 2018; Kato *et al*., 2020; Debracque *et al*., 2022; Hayashi *et al*., 2022, 2022). Our study, by demonstrating cortical activation in predefined regions associated with social decision-making, particularly regions across various social scenarios, confirm their pivotal role in primate social cognition, and makes a significant contribution to this nascent field of social neuroscience. This awake, head-free paradigm overcomes critical limitations of anaesthesia-dependent studies (Yamada *et al*., 2018; Kato *et al*., 2020), revealing nuanced social context effects that passive designs cannot capture (Testard *et al*., 2024).

We identified an early haemodynamic signature characterised by reduced oxygenated haemoglobin (HbO) and increased deoxygenated haemoglobin (HbR) during the initial 3 seconds of coaction, envy, altruism, and competition tasks. This “initial dip” suggests rapid oxygen consumption in socially engaged regions (cf. Rizzolatti & Sinigaglia, 2010), potentially reflecting the immediate cognitive demands of processing complex social interactions.

The subsequent GLM analysis showed condition-dependent variations in sustained neural engagement: altruism, competition, and cooperation elicited stronger haemodynamic responses (higher B₁ values), consistent with the involvement of frontal regions in goal-directed social behaviours.

Conversely, the attenuated responses during coaction tasks may indicate either cortical inhibitory processes or greater subcortical involvement during these interactive contexts.

The observed dlPFC-TPJ-STS activation patterns mirror those reported in human social cognition studies (Donaldson *et al*., 2015), supporting the conservation of these neural circuits across primates. Notably, the differential haemodynamic profiles between social conditions (e.g., strong initial dip in envy versus sustained response in cooperation) suggest distinct neural computation strategies for various social challenges. These findings establish an important foundation for using macaque models to study the neurobiology of social behaviours and their potential disruptions in psychiatric disorders.

The observed patterns of prefrontal and temporoparietal activation in macaques during dyadic social interactions exhibit striking convergence with human fNIRS studies, underscoring conserved neural substrates for social cognition across primates. Our macaque paradigm, though not employing hyper-scanning, similarly captures dyadic social dynamics through its face-to-face design and could be adapted for future inter-brain studies in primates. Notably, the prefrontal cortex (PFC) and temporoparietal regions in human emerges as a pivotal hub for coordinating joint actions in both species. Human hyper-scanning research demonstrates that inter-brain synchrony in the dlPFC predicts cooperative success during joint tasks like gambling (Zhang *et al*., 2017), mirroring our findings of heightened dlPFC engagement during macaque coaction. This alignment suggests an evolutionarily preserved mechanism for action coordination, where the dlPFC integrates cognitive control and social monitoring to facilitate interactive behaviours. Further reinforcing this parallel, studies of mother-child dyads reveal dlPFC synchrony during cooperative tasks (Bizzego *et al*., 2022), hinting that such neural coupling may generalise to diverse social bonds.

Very few studies have also shown different haemodynamic responding patterns to positive and negative emotions within the PFC and STS (Yükselen *et al*., 2023; Tang *et al*., 2025), or positive emotion clusters in frontal neural activities (Hu *et al*., 2019), and the fact that this haemodynamic response can disrupt in social disorders like schizophrenia (Wang *et al*., 2023). Intriguingly, sex-specific modulations of dlPFC coherence in humans—such as heightened inter-brain synchrony during risky decision-making, which raise compelling questions about whether similar sociodemographic factors influence macaque neural dynamics, a promising avenue for future cross-species investigation.

Temporoparietal regions, particularly the temporoparietal junction (TPJ), further illustrate this cross-species homology. Human studies consistently implicate the TPJ in mentalising, empathy, and emotional synchrony, and involving in bidirectional information flow between interacting brains (Goelman *et al*., 2019), resonating with our macaque data during empathy-driven interactions. For instance, TPJ activity in humans correlates with emotional valence alignment during communication, with heightened haemodynamic responses to negative arousal (Yükselen *et al*., 2023). Similarly, sex differences in TPJ coherence—such as increased inter-brain synchrony in females during high-risk social tasks—suggest its role in empathic processing and nonverbal cue integration, paralleling our observations of TPJ modulation in macaques (Zhang *et al*., 2017).

The superior temporal sulcus (STS) serves as a key node for social perception, with human studies implicating it in emotional communication and bidirectional signalling during joint attention interactions, as demonstrated by fMRI research (Goelman *et al*., 2019). fNIRS evidence further highlights its sensitivity to social context, showing that haemodynamic responses in the STS are enhanced by social inclusion during biological motion processing (Bolling *et al*., 2013; Tang *et al*., 2025).

These findings demonstrate important consistencies and opportunities in comparative social neuroscience research. The robust involvement of dlPFC and TPJ in both human and macaque studies confirms their essential role in cooperative behaviour and empathy processing, aligning with established human fNIRS literature on joint action and mentalising. Human studies of competition present more complex patterns, with some reports showing frontal coherence changes while others indicate global network integration. This variability suggests that competitive neural signatures may depend on specific task parameters, an area where controlled primate studies could provide valuable clarification. Notably, existing human fNIRS research has not directly examined envy responses, making our macaque findings particularly significant for understanding this evolutionarily conserved social emotion. The current results establish a foundation for investigating how basic social emotions engage shared neural circuits, particularly in frontal regions associated with social evaluation processes.

These neural parallels gain further significance from human hyper-scanning research, which links interpersonal neural alignment in frontal regions to cooperation success. Though our study did not employ hyper-scanning, the face-to-face macaque paradigm nevertheless captures analogous dyadic dynamics through synchronised behavioural and physiological measurements, supporting the hypothesis that primate social cognition operates through a core set of interacting neural systems. Moreover, the successful detection of socially-driven activity in macaques expands fNIRS applications beyond motor functions, demonstrating its efficacy for probing higher-level cognition in non-human primates.

### 3.6. Limitations

Several important limitations should be considered when interpreting these findings. First, we observed individual differences in decision-making patterns (like humans, which show great variation in behaviour and personality, De Petrillo & Rosati, 2021), and our small sample size (n=6) and use of same-age, same-sex dyads limit our ability to examine how factors like sex differences, familiarity, or developmental stages influence social cognition. Second, we did not observe a familiarity effect, likely because all six monkeys were raised together from childhood and housed in the same facility room. While we kept pairs consistent during the 3-year study, their pre-existing social bonds may have masked familiarity-related neural responses. Future work could test novel pairings (e.g., monkeys from separate facility rooms) or even cross-species dyads (e.g., rhesus vs. long-tailed macaques) to clarify this.

Third, the observed behavioural variation, particularly lower altruism responses than those reported in macaques (Ballesta & Duhamel, 2015), suggests potential genetic, neurochemical (e.g., oxytocin), training, or early-life differences that future studies should explore. However, the majority of existing studies (Chang *et al*., 2015; Sallet *et al*., 2021) indicate that macaques show concern for others only when the choice involves both self and other rewards, and no subjects chose altruistically when selecting between self-only and other-only rewards. Lastly, fNIRS’s ∼2.5 cm penetration depth limits assessment of subcortical areas (e.g., amygdala, cingulate) involved in social-emotional processing. Combining fNIRS with other modalities (e.g., PET/MRI) could address these spatial and temporal resolution constraints.

### 3.7. Summary and Future Directions

Building on these findings, several promising research avenues emerge. First, pharmacological interventions using oxytocin could be employed in dyadic tasks, particularly using monkey models to evaluate its potential as a therapeutic intervention approach for social deficiencies (Zarei *et al*., 2020), while examining how its effects vary across different social contexts (Zarei *et al*., 2019a). Second, comparative studies across primate species (Wang & Liu, 2023) could elucidate both conserved and species-specific neural mechanisms underlying social decision-making, as it is suggested that different primate species shows different level of social cognitive abilities.

Methodologically, combining fNIRS with electrophysiological recordings would provide complementary temporal resolution to haemodynamic measures. Including analytical approaches, such as diffuse optical tomography and effective connectivity analyses, could offer more comprehensive insights into the neural mechanisms driving observed haemodynamic changes. Finally, this paradigm could be adapted to model neuro-developmental conditions like autism or schizophrenia (Testard *et al*., 2024), investigating how social contexts differentially engage key regions such as the dlPFC, TPJ, STS, and other related brain area to social interaction and in different conditions, or scrutinise the role in theory of mind (Wang *et al*., 2025) while examining broader network dysfunction across social cognition domains.

## 4. Materials and Methods

### 4.1. Subjects and Housing

Six male long-tailed macaques (*Macaca fascicularis*; age: 3 years at training onset, weight: 4 ± 0.5 kg) participated in this study. All monkeys were naïve to cognitive tasks prior to the experiment and were pair-housed in standard primate cages under a 12:12-hour light-dark cycle. Subjects were reared together from infancy, allowing clear establishment of dominance hierarchies.

To maintain monkeys’ motivation to perform the tasks, training and testing were conducted prior to their daily meal. The monkeys received rewards while performing the tasks and were provided with their most part of their daily fresh fruit and vegetable immediately after daily testing to encourage monkey to come to the lab for testing.

In the evening, they were also given a reduced amount of food and water. We conducted our experiments in highly controlled conditions, prioritising the parsimonious use of animals. The weight and welfare of the animals were closely monitored by the animal facility staff, veterinarians, and lab scientists. All procedures complied with the NIH Guide for the Care and Use of Laboratory Animals and the Chinese National Guidelines for Animal Research, and were approved by the ION, CAS Animal Ethics Committee.

The study used a non-invasive touchscreen-based paradigm, eye-tracking, and head-free fNIRS system, requiring no surgical implantation. Social cognitive tasks were conducted in a controlled environment to minimise stress.

### 4.2. Experimental setup

The computerised testing station comprised two interconnected systems: an Ubuntu 22.04 workstation and a Windows 11 workstation. The Ubuntu system controlled behavioural task presentation using the Opticka experiment manager (Andolina, 2023) with Psychtoolbox (PTB) (Kleiner *et al*., 2007) in MATLAB 2022a, while simultaneously recording video from three behaviour cameras. The Windows system operated the two eye-tracking setups: a head-restricted system using iRecHS2 software (Matsuda *et al*., 2017) and a head-free system using Pupil Capture/Player (Pupil Labs). These systems communicated via a local Ethernet hub using TCP and UDP protocols to ensure synchronisation.

Stimuli were displayed on a 55-inch Xiaomi OLED transparent touchscreen (LG panel) with a resolution of 1920 × 1080 pixels, a 120 Hz refresh rate, a 15,000,000:1 contrast ratio, and a 1 ms response time. The display featured dual-sided capacitive touch input (120 Hz temporal resolution) with antiglare and tempered glass, enabling two monkeys to interact face-to-face while viewing identical stimuli (fig. 1). The subject positions on either side of the transparent OLED (tOLED), partner assignments, and the experimental conditions were pseudorandomly changed across sessions.

Pairs of macaques were seated face-to-face and performed a paired foraging-like search task by touching designated visual targets on the shared screen in different conditions to receive food rewards (dustless precision pellets, Bio-Serv). Behavioural measures included accuracy (correct responses) and reaction time (touch latency), synchronised with eye-tracking data.

The monkeys were seated at an angle to the transparent OLED screen (**Fig. 1**) for two important reasons. First, this angular placement prevented facial reflections from appearing directly on the screen. Instead, any reflections were redirected to the sides where anti-glare glass panels were installed. A direct 90-degree seating arrangement might cause slight distracting facial reflections in the central display area. Second, the angled positioning created distinct interactive zones on the touchscreen (**Fig. 11**). While a 90-degree face-to-face arrangement would have given both monkeys equal access to identical screen areas, our angled monkey location allowed for more flexible task designs. We could either present shared targets in a central mutually-accessible zone, or display separate stimulus sets in distinct lateral regions, with each monkey having exclusive access to their designated area while remaining unable to reach their partner’s stimuli.

**Figure 11.**
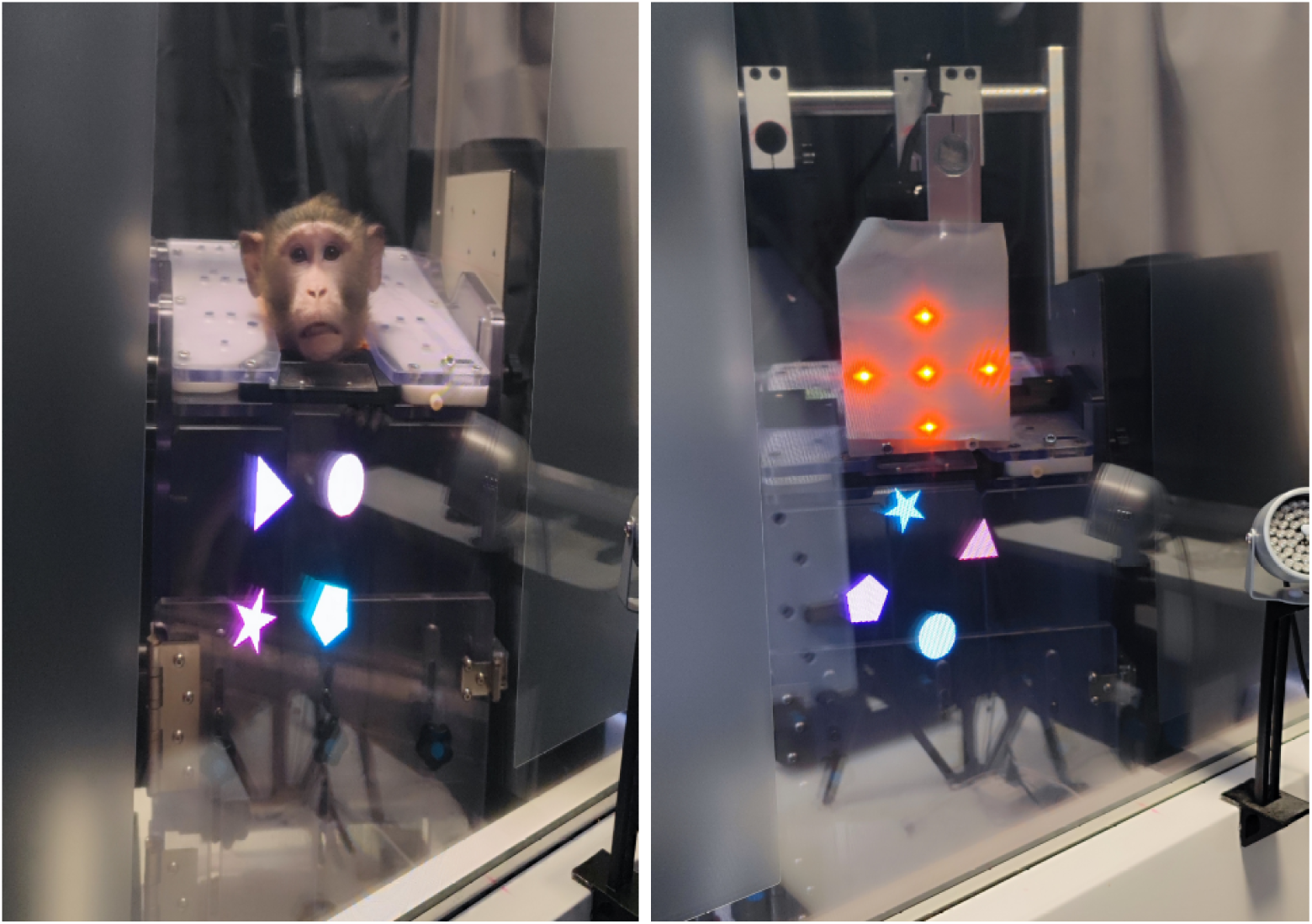
Visual Identification of the other with stimulus presentation. (Left) Monkeys interacting through the transparent touchscreen display which presented stimuli. (Right) Red LED light boards used for five-point eye-tracker calibration on the other side prior to sessions (displayed simultaneously with stimuli to establish spatial reference points). As shown, stimuli were positioned randomly rotated around a virtual circle with randomised colours. Anti-glare film prevented monkeys from seeing their own reflections on the screen.

### 4.3. Training Procedure and Tasks

We started training monkeys to precisely touch the target to receive a reward and then prepare for the main task in their home cage, using an in-cage wireless automated touchscreen system for their initial training (github.com/CogPlatform/CageLab). All monkeys had to first reach a stable performance level of 80% in each stage of training under the in-cage training system. Then, they moved to the lab for system familiarisation and investigation of social interaction in the main social task and sat in a transparent plexiglass monkey chair on either side of the tOLED, so monkeys could see each other’s face and body, as well as the target on the shared screen.

However, when we brought the monkeys into the room, we covered the transparent tOLED and then, for calibration (We calibrated for the partner monkey’s facial region), there was a light board (**Fig. 11**) in front of the other monkey’s face for calibration. This ensured that the two monkeys couldn’t see each other before the main task began. The testing room had a dim light, so that subjects could clearly see each other’s face and the stimuli were clear on the shared tOLED from both sides. We used fluid reward for eye-tracking calibration and pellet rewards for the main tasks. This was because fluid rewards on one side might misleading their partner’s judgment about the partner’s reward reception, as monkeys’ mouths were usually on the water tube and they might lick or suck the water tube even without a reward. We used food pellets (190-mg food pellet; Dustless Precision Pellets for Primates, Bio-Serv Co., USA) as the reward to provide a clear event for both subjects participating in a social interaction.

We used two types of tasks with eye-tracking: reward-cued shape tasks with iRecHS2 eye-tracking and a ping-pong set game with head-free Pupil Core eye-tracking, as we aimed to have tasks with head-free fNIRS recording and a cooperative context. Both tasks’ design included a range of social contexts:

#### 4.3.1. Task 1: Reward-Cued Object Categorisation Task

Our main tasks included four geometric stimuli that were visible for both participants. Each shape had a different reward contingency. Star (★Self): touching the star led to receiving a reward immediately by the subject. Circle (●None): no reward; neither the self nor the partner received a reward. Polygon (⬣Both): touching this stimulus both subjects received a reward; first the partner received a reward immediately, and after a 1s delay, the subject received a reward. Triangle (▲Other): The subject didn’t receive any reward, but the monkey on the other side would receive the reward. The colour and the location of each stimulus on the screen were changed randomly at each trial, preventing the monkey from learning to touch a specific stimulus at the same place or by colour without attention to the task.

The stimuli were presented to animals for 3s in each trial. Touching each stimulus resulted in a auditory cue for different outcome and the disappearance of all stimuli. If the touch was correct, a food pellet was dispensed by a computer-controlled pellet dispenser. The next trial began after 3 seconds (the inter-trial interval). Experimenters monitored the animals’ behaviour and task progression from a control room through the three behavioural cameras positioned within the testing cubicle.

In the control group, all four stimuli were presented to a single monkey alone in the testing room. Rewards were delivered based on the touched stimulus. If the monkey selected a target that (also) rewarded the partner (e.g., ⬣Both or ▴Other), the partner’s reward was collected in a transparent glass bowl, allowing the acting monkey to observe the pellet delivery on the other side.

In the presence of a partner, the task was performed with a controlled set of possible social interactions:

1. Audience Context: The subject performed the task while the partner just sat on the other side. The partner did not receive any reward and remained slightly farther from the screen to avoid touching it. All four stimuli were presented as in the control group, and the partner received no reward. Rewards for ⬣Both or ▴Other were collected in a transparent glass on the partner’s side.
2. Co-action Context: Two sets of stimuli were presented (each monkey could only reach their own set). Both monkeys actively performed the task and received a reward when touching the proper answer, regardless of the other monkey’s answer (only ★Self and ●None stimuli were presented).
3. Envy Context: A single subject performed the test with the ⬣Both and ●None stimuli. Touching ⬣Both rewarded both monkeys: the partner immediately, and the actor monkey after a 1s delay. Different auditory tones accompanied each reward delivery.
4. Altruism Context: ▲Other and ●None stimuli were presented. The partner received the reward for the subject’s correct response, requiring the subject to act just for the other monkey’s benefit. To prevent task abandonment, we included occasional Star (Self) trials. The number of Triangle touches was recorded.
5. Competition Context: A single set of stimuli was presented, and both subjects competed to respond faster. Only the faster monkey received a reward with ★Self and ●None stimuli presented.

#### 4.3.2. Task 2: Ping Pong Tasks

In the Ping-Pong Game, we sought to simulate a more natural cooperative experience by incorporating a single ball both subjects manipulated and which obeyed physical laws using a physics engine, allowing a virtual ball to behave according to realistic characteristics such as gravity, speed, air resistance etc.

The first subject dragged a virtual ball from one side of the screen to the other side, and the other monkey should return it back, enabling both to receive a reward. The Ping-Pong cooperation task required subjects to engage with their partners as integral participants in the trial. They had limited time to complete the task, and if either monkey failed to participate, neither received a reward. This design necessitated mutual attention between partners.

Like the reward-cued shape task, Ping-Pong also included all scenarios structured in addition to the Ping-Pong cooperation task: 1. Control Group: In this scenario, the subject must drag a virtual ball towards the other side of the screen, ensuring the ball crosses the central line to receive rewards. 2. Audience Effect: Here, the subject performs the task while the partner simply observes from the other side of the screen, without receiving any rewards. The partner is positioned slightly away from the screen to avoid touching the screen. 3. Co-action Effect: Both monkeys actively participate in the task (two balls, one for each monkey), with rewards given for successfully dragging their virtual balls to the central line, independent of each other’s performance. 4. Envy Effect: In this scenario, a single subject drags a virtual ball through the central line, and both monkeys receive rewards. The partner is rewarded immediately after the correct response, while the subject receives their reward after a one-second delay. Different auditory tones accompany the reward delivery for each monkey. 5. Altruism Effect: In this case, the partner receives the reward while the subject drags the ball. The subject performs the task solely for the benefit of the other monkey, without receiving any reward themselves. 6. Competition Effect: In this scenario, both monkeys strive to perform the task sooner, and only the monkey who reaches their ball to the central line first, receives a reward, and the trial finishes, and the other partner’s ball also disappears. A flash of two different background colours appears with specific sounds for back and front, to emphasise which monkey received the reward. 7. pingpong cooperation task which we mentioned at the beginning.

### 4.4. Behavioural Recording and Eye-tracking

Eye-tracking is a valuable tool especially in studying nonverbal subjects like animals and human infants, enabling researchers to gauge their knowledge and surprise levels through measures such as pupillometry and looking time. Similar to human studies (Sim & Xu, 2019; Zarei *et al*., 2022), other animals exhibit pupil dilation and longer looking times when they are surprised by a situation’s outcome (Machado & Nelson, 2011; Kuraoka & Nakamura, 2022). This method has been applied across various nonverbal populations, including human infants and different primate and nonhuman animal species. Additionally, eye-tracking provides insights into ecologically relevant responses and potential arousal levels by measuring pupil diameter. By employing eye-tracking techniques, researchers can delve deeper into macaque behaviour, uncovering new insights into social decision-making tasks and better understanding the connection between visual input and behavioural responses.

We also used three USB3 video cameras (Sony IMX290; 1920×1080; 60 fps) for behavioural recording: one camera on each side of the setup to record the monkeys’ faces and one camera above the setup for action monitoring. Three cameras for behaviour recording were monitored and synchronised using OBS Studio. For our study, we used two types of eye-trackers: head-fixed iRecHS2 and head-free Pupil Core systems.

#### 4.4.1. iRecHS2 Eye Tracking

We utilised two Chameleon3 USB3.1 industrial cameras (Model: CM3-U3-13Y3M; 1.3 MP, 500 FPS, CS-mount, USB 3), two Fujinon C-mount lenses (Model: Hf16HA-1S, f/1.4, 16 mm), an IR filter, and an infrared fill light illuminator (850 nm, 60 mm). Before each session, we calibrated each monkey’s eyes using Flycapture2 SDK, iRecHS2 software and a MATLAB script. The two eye-tracking cameras were synchronised with the presentation system via TCP/UDP to receive triggers at the start of each session and trial.

The noninvasive head-restraint method for calibration used a light-weight custom 3D-printed helmet based on a structural MRI head model from each monkey, tethered to the monkey chair. The iRec system used infrared illumination and cameras to detect pupil/corneal reflections, integrating these with an internal model to compute gaze data (sampled at 500 Hz). Gaze data were synchronised with the tOLED display. A five-point calibration required fixation within 0.5° visual angle for 75 ms. The eye tracker was mounted on the monkey’s chair, positioned frontolaterally at 10 cm distance to avoid visual field occlusion. We performed eye-tracking calibration and validation for each monkey before each session. To calibrate, we placed a light board (Fig. 2) in front of the other monkey’s face, featuring five LEDs (central and peripheral). During calibration, monkeys could not see each other. We defined the region of interest (ROI) as a 6 cm diameter circle covering the monkey’s head. We evaluated how social interactions across conditions affected gaze patterns toward the ROI and whether dominance influenced looking time or ROI fixation duration.

#### 4.4.2. Pupil Core Head-free Eye-Tracking System

For the second phase, we deployed two Pupil Core headsets (Pupil Labs GmbH) modified with a customised 3D printed helmet for NHP use simultaneously on both sides (scene camera: 1080p–480p @ 30–120 Hz; eye cameras: 2 × IR, 192 × 192 px @ 200 Hz; 2D/3D pupil and gaze measurements), with a nominal precision of 1° visual angle. 0.60° accuracy; 0.02° precision. The head-free eye-tracking system allowed monkeys to move naturally during cognitive tasks while maintaining measurement accuracy, thereby preserving more ecologically valid social interactions between paired subjects. This configuration captured precise visual attention data, enabling detailed analysis of gaze patterns in response to both social partners and experimental stimuli. For our monkey studies, we implemented the Pupil Core eye-tracking system (Pupil Labs GmbH) with custom-designed frames individually fitted to each subject’s head morphology (**Supp. Fig. 2**). To maximise naturalistic behaviour, the system permitted full head and hand mobility. We mounted a plexiglass panel featuring a small access port to each chair, allowing reward retrieval without interfering with recording equipment.

Calibration and validation procedures required modifications to the Opticka experimental control code. The system employed two synchronised eye-trackers positioned on opposite sides of the experimental setup. Due to the computationally intensive nature of the Pupil Capture software, we distributed processing across two computers: one running MATLAB/Opticka with a single Pupil Capture instance, and a second dedicated computer operating the complementary eye-tracker and Pupil Capture software. Synchronisation between systems was maintained through a local Ethernet hub using TCP and UDP protocols, ensuring simultaneous operation initiation. Each Pupil Core eye-tracking unit consisted of a scene camera supporting multiple resolution and frequency configurations (including 1080p at 30 Hz, 720p at 60 Hz, and 480p at 120 Hz), paired with dual infrared eye cameras capturing at 192 × 192 pixel resolution and 200 Hz sampling rate. The system provided comprehensive 2D and 3D pupil and gaze vector measurements, achieving 0.60° accuracy following calibration, 0.02° precision, and 1° visual angle resolution for optimal spatial tracking fidelity.

### 4.5. Functional Near-Infrared Spectroscopy (fNIRS)

A key advantage of fNIRS for primate research is its non-invasiveness, portability, and relatively low susceptibility to motion artefacts compared to techniques like fMRI and PET. This allows for brain activity measurements under conditions with fewer body movement constraints, facilitating investigations of more naturalistic behaviours. In addition to better assessing innate or behaviourally relevant abilities, we aimed to explore techniques that have transformed behavioural testing by providing a richer context for behavioural and cognitive analysis. We employed the NirSmartⅡ-3000A fNIRS system (Danyang Huichuang Medical Equipment Co., Ltd., China) to monitor haemodynamic responses in social decision-making tasks (Jiao *et al*., 2024; Liu *et al*., 2025; Zhang *et al*., 2025). The system targeted key cortical regions involved in social cognition, including the dorsolateral prefrontal cortex (dlPFC) and parietal areas (superior temporal sulcus [STS] and temporoparietal junction [TPJ]) in both hemispheres.

#### System Specifications

The fNIRS system utilised near-infrared light-emitting diodes (LEDs; 730 nm and 850 nm wavelengths) paired with avalanche photodiode (APD) detectors, sampling at 11 Hz. Our optode configuration consisted of 16 optodes (8 sources, 8 detectors) arranged in a custom head cap with 1.6 cm source-detector separation distance. Through empirical testing of various distances (1.6 cm, 2.1 cm, and 2.6 cm), we determined 1.6 cm provided optimal coverage of our regions of interest while maintaining signal quality for macaque neuroimaging.

#### Head cap design

Developing effective fNIRS head caps for macaques required overcoming several anatomical challenges. Unlike humans, macaques exhibit significant variation in cranial morphology and lack a defined forehead structure, making stable optode placement particularly difficult during head-free experiments. These factors are especially problematic given that most commercial optode arrays are designed for human neuroimaging.

To ensure accurate and stable positioning over our target cortical regions, we created individualised caps based on each subject’s MRI scans and 3D-printed cranial models (**Fig. 12**). The custom design accommodated the subjects’ neuroanatomical variations while maintaining the critical 1.6 cm source-detector separation. Special attention was given to the cap’s mechanical stability to prevent movement artefacts during task performance, incorporating features to compensate for the absent forehead structure. This customised approach allowed reliable monitoring of haemodynamic responses (ΔHbO, ΔHbR, and ΔHbT) throughout behavioural tasks, providing robust data for analysing neural correlates of social decision-making.

**Figure 12.**
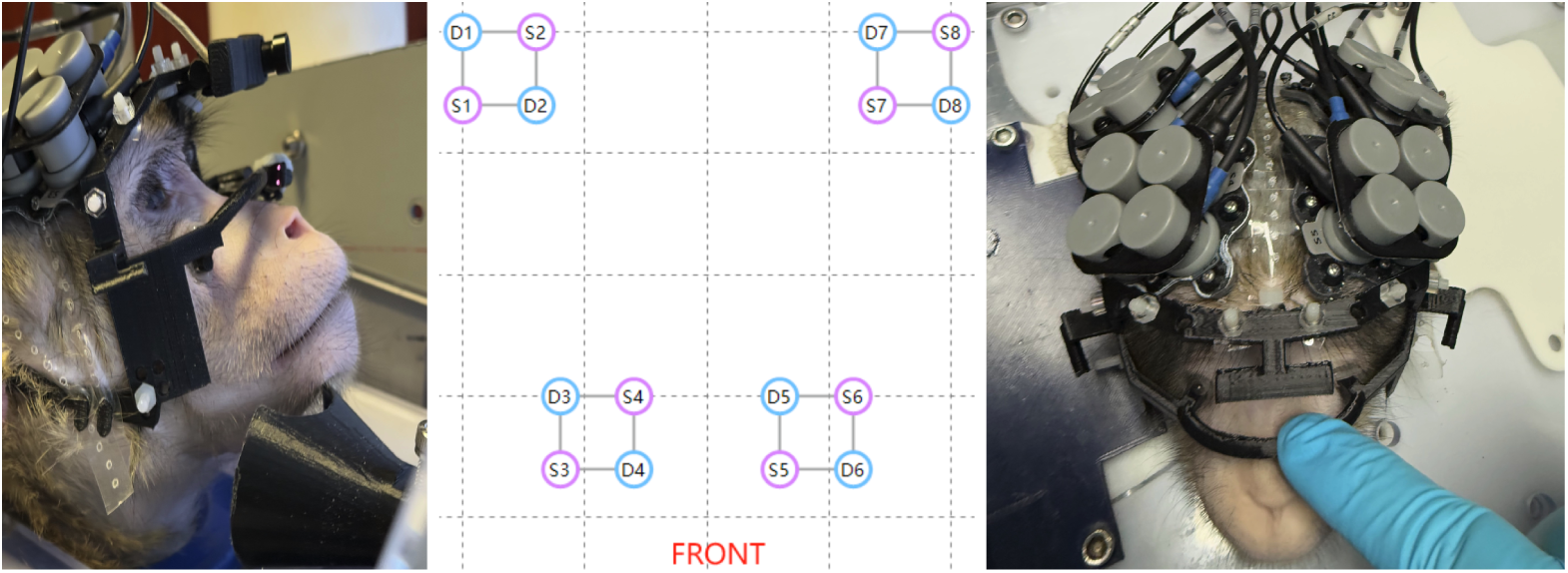
Dual-modal neurobehavioral recording setup. A macaque wearing the custom fNIRS cap and Pupil Core head-free eye-tracking system during task performance, enabling simultaneous measurement of cortical haemodynamics and gaze patterns. (Centre) Optode arrangement on the macaque’s head, showing source-detector placement over target regions of interest (dlPFC, STS/TPJ).

#### Combination of head-free Eye-tracking and fNIRS

The integration of head-free eye-tracking and fNIRS represents a significant advancement in the study of cognitive processes in non-human primates. By combining these two technologies, we aim to gain a more comprehensive understanding of the emotional states and neural mechanisms underlying social decision-making and interactions more naturally. While eye-tracking provides real-time data on visual attention and social engagement, fNIRS enables us to correlate these behaviours with specific neural responses. This dual approach not only enhances our understanding of individual decision-making in social contexts but also sheds light on the neural underpinnings of complex cognitive behaviours, establishing a robust framework for future research that contributes to our understanding of social cognition and its related neural mechanisms in non-human primates and explores the motivational and affective underpinnings of prosocial behaviour through social decision-making.

#### fNIRS Data Preprocessing

Raw NIRS data were processed using NirSpark software (v1.7.5; Huichang, China). Optical density data were converted to oxygenated (ΔHbO) and deoxygenated haemoglobin (ΔHbR) concentration changes via the modified Beer-Lambert law, applying differential path-length factors (DPF) of 5 (730 nm) and 6 (850 nm) to account for wavelength-specific light scattering in macaque tissue.

#### Quality Control, Filtering, and Processing

Channels were excluded based on three criteria: (1) signal-to-noise ratio (SNR) < 2, (2) optical intensity outside the 0.5–1000 range, or (3) coefficient of variation (standard deviation/mean) < 2. Motion artefacts were corrected using threshold-based filtering (amplitude threshold = 0.5; standard deviation threshold = 6). Physiological noise was attenuated via bandpass filtering (0.01–0.2 Hz), followed by second-order detrending to eliminate signal drift. Principal component analysis (PCA) was applied to remove the first component corresponding to the largest eigenvector, effectively reducing systemic physiological contamination. The continuous wavelet transform was then employed to isolate neurophysiological haemodynamic responses within the 0.01–0.2 Hz frequency band, which captures both resting-state fluctuations and task-evoked activity.

#### Frequency Spectrum

To confirm the physiological origin of our fNIRS signals, we performed spectral validation in both anaesthetised and awake states. Under anaesthesia (heart rate: 2-2.5 Hz), simultaneous fNIRS-ECG recordings showed clear cardiac peaks in the power spectrum (**Supp. Fig. 3**, top). In awake monkeys (heart rate: ∼3.5 Hz), these peaks shifted to higher frequencies while maintaining detectable amplitudes (**Supp. Fig. 3**, bottom). This confirmed our system’s sensitivity to cardiopulmonary signals while demonstrating their spectral separation from the 0.01-0.2 Hz neurogenic haemodynamic band. Channel stability was quantified using the coefficient of variation (CV = SD/mean × 100). Following NirSpark software’s quality criteria, channels exceeding CV > 20 were excluded as outliers.

### 4.6. Statistical Analysis

All analyses were performed using JASP (JASP Team, 2024). We first assessed data normality using Shapiro-Wilk tests and homogeneity of variance with Levene’s tests. For repeated measures data meeting parametric assumptions, we conducted repeated-measures ANOVAs, verifying sphericity with Mauchly’s test and applying Greenhouse-Geisser corrections when violations occurred. Significant main effects were followed by Bonferroni-Holm corrected post-hoc tests. We complemented these analyses with effect size measures including Cohen’s d (interpreted as: >0.2 small, >0.5 medium, >0.8 large) and partial omega-squared (ω²ₚ: >0.01 small, >0.06 medium, >0.14 large). All data are expressed as mean ± SEM unless otherwise noted, with statistical significance threshold set at p < 0.05. Where applicable, we additionally reported rank-biserial correlations for non-parametric effect sizes.

For non-normal data or when assumptions were violated, we employed non-parametric alternatives. The Friedman test served as our primary non-parametric repeated measures analysis instead of repeated measure ANOVA, with Kendall’s W for effect size interpretation (0.1 small, 0.3 medium, 0.5 large). As Friedman’s test cannot accommodate between-subjects factors, we supplemented it with Mann-Whitney U tests (Bonferroni-Holm corrected) for between-group comparisons (like dominancy) when data showed acceptable skewness and kurtosis (<2). Significant Friedman results were followed by Conover post-hoc tests with appropriate corrections.

To further strengthen our inferences, we computed Bayes factors (BF₁₀) using default JASP priors (Rouder, 2012). These quantified the relative evidence for alternative versus null hypotheses, with BF > 3 indicating substantial support for the alternative and BF < 0.5 supporting the null. This Bayesian approach proved particularly valuable for testing complex hypotheses involving equality and order constraints, while remaining interpretable as evidence weights for competing theories (Dienes, 2014; Mulder *et al*., 2021).

## 5. Supplemental Information

**Supplementary Figure 1.**
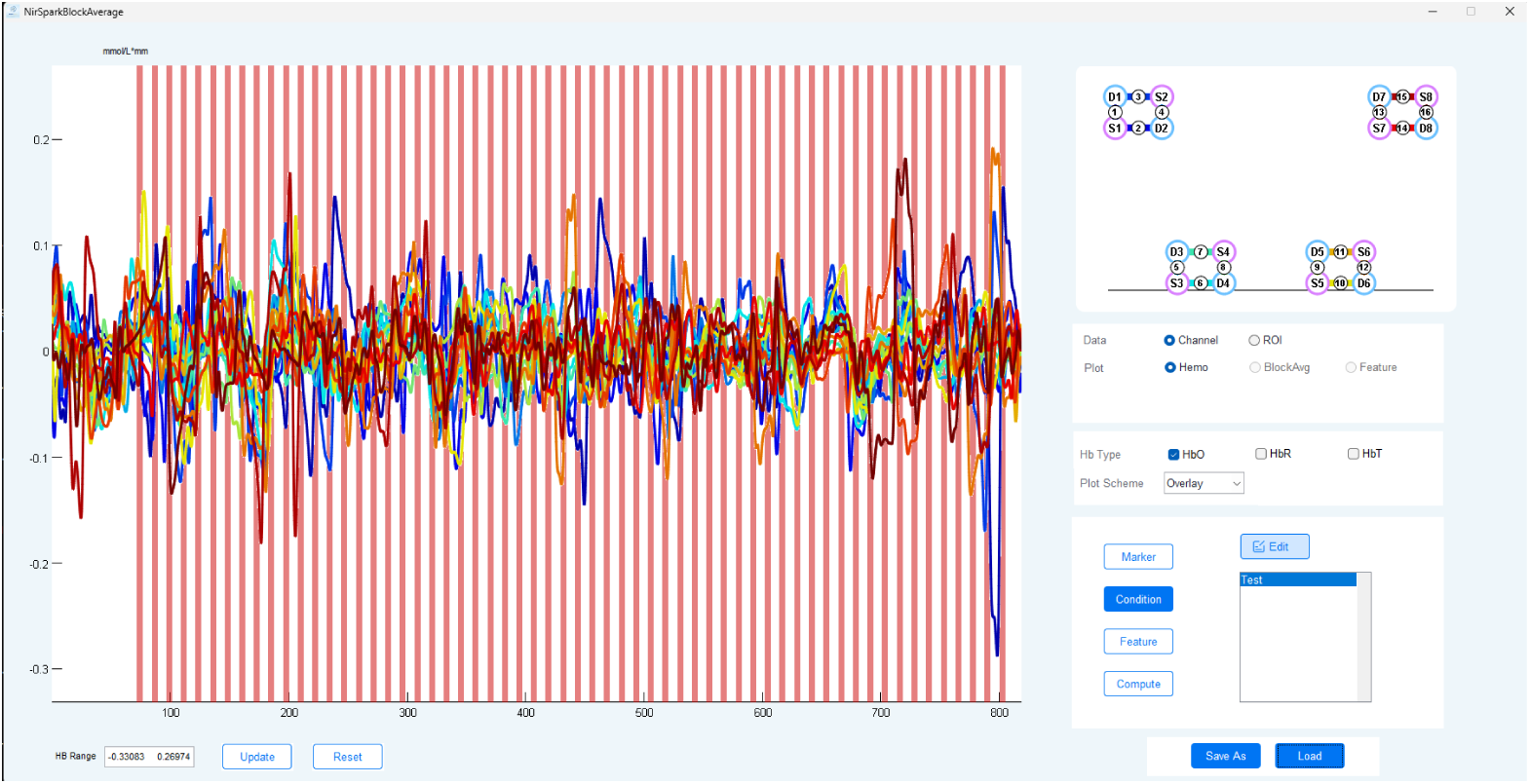
Haemodynamic responses across channels, aligned to task markers in a session. Overlay of HbO concentration changes from all recorded channels aligned with task timing markers. Red shaded areas indicate 3-second active task periods; intervening white spaces represent 4-second inter-trial rest intervals.

**Supplementary Figure 2.**
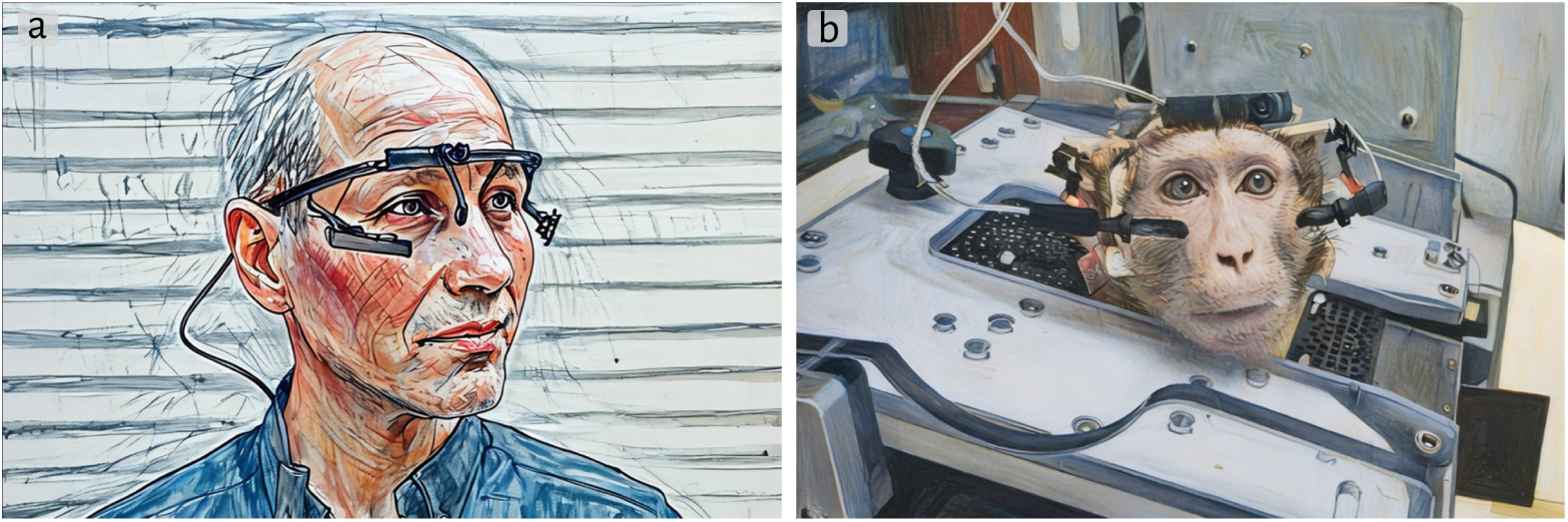
Non-invasive head-free eyetracking in non-human primates. (**a**) Head-free eye-tracking system from Pupil Labs. (**b**) 3D-printed a custom holder (see also Fig. 12) to adapt the Pupil core world and eye cameras configured for head-free monkey use.

**Supplementary Figure 3.**
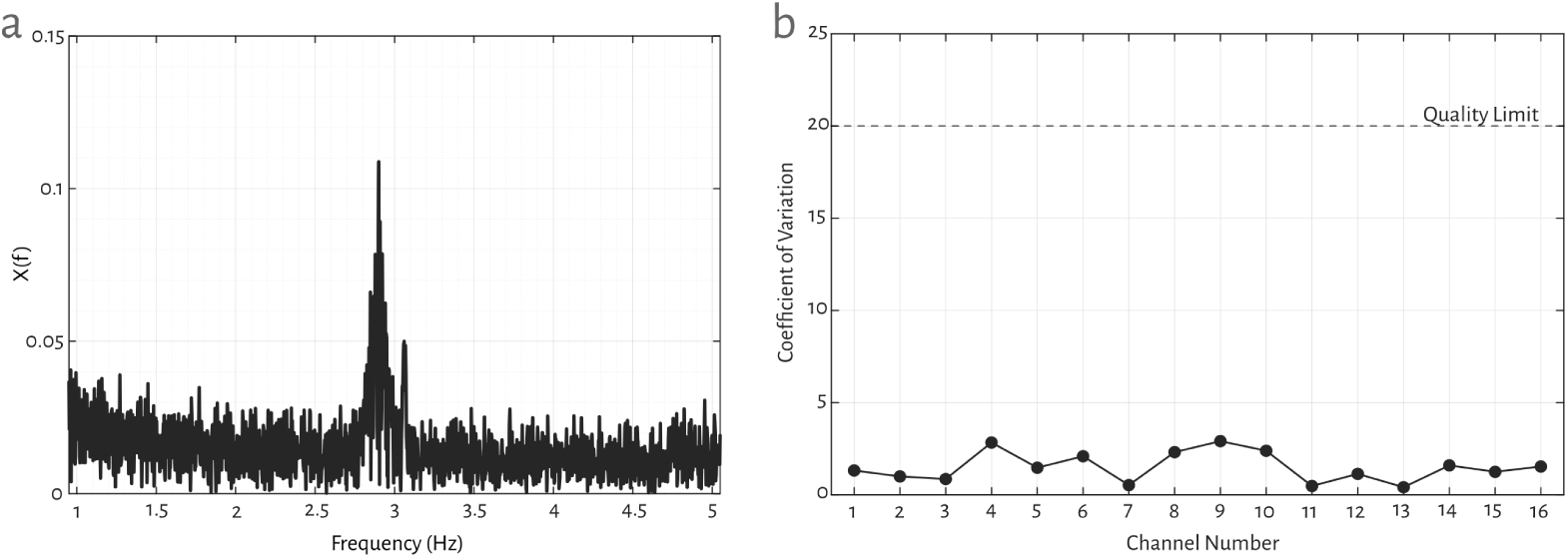
Validation of cardiac signals in fNIRS recordings and coefficient of variation (CV). (**a**) Power spectrum from a monkey showing distinct cardiac peaks (∼2.8 Hz). (**b**) Channel-wise coefficient of variation (CV) across 16 fNIRS channels during task performance.

## 6. Footnotes

### Footnotes Funding

This work was supported by the following grants: Shanghai Municipal Science and Technology Major Project 2018SHZDZX05 (to I.M.A.), National Natural Science Foundation of China grants 32070992 (to I.M.A.) and 32150410370 (to S.Z.).

### Author Contributions

**Shahab A. Zarei**: Conceptualisation, Experimental section, Data collection and analysis, Investigation, Methodology, Writing original draft, review and editing; **Jiahao Tu**: Animal management, experimental technical development; **Xiaochun Wang**: experimental engineering support; **Ian Max Andolina**: Supervision, Conceptualisation, Writing original draft, review and editing, Visualisation, Data monitoring and analysis and curation, review and editing, Visualisation. All authors read and approved the final manuscript.

### Declaration of Conflicting Interests

The authors declare that there is no conflict of interest.

## Notes

### Competing Interest Statement

The authors have declared no competing interest.

